# The Mayo Clinic Salivary Tissue-Organoid Biobanking: A Resource for Salivary Regeneration Research

**DOI:** 10.1101/2024.02.23.581761

**Authors:** Syed Mohammed Musheer Aalam, Ana Rita Varela, Aalim Khaderi, Ronsard J Mondesir, Dong-Gi Mun, Andrew Ding, Isabelle M.A. Lombaert, Rob P. Coppes, Chitra Priya Emperumal, Akhilesh Pandey, Jeffrey R. Janus, Nagarajan Kannan

## Abstract

The salivary gland (SG) is an essential organ that secretes saliva, which supports versatile oral function throughout life, and is maintained by elusive epithelial stem and progenitor cells (SGSPC). Unfortunately, aging, drugs, autoimmune disorders, and cancer treatments can lead to salivary dysfunction and associated health consequences. Despite many ongoing therapeutic efforts to mediate those conditions, investigating human SGSPC is challenging due to lack of standardized tissue collection, limited tissue access, and inadequate purification methods. Herein, we established a diverse and clinically annotated salivary regenerative biobanking at the Mayo Clinic, optimizing viable salivary cell isolation and clonal assays in both 2D and 3D-matrigel growth environments. Our analysis identified ductal epithelial cells in vitro enriched with SGSPC expressing the CD24/EpCAM/CD49f+ and PSMA-phenotype. We identified PSMA expression as a reliable SGSPC differentiation marker. Moreover, we identified progenitor cell types with shared phenotypes exhibiting three distinct clonal patterns of salivary differentiation in a 2D environment. Leveraging innovative label-free unbiased LC-MS/MS-based single-cell proteomics, we identified 819 proteins across 71 single cell proteome datasets from purified progenitor-enriched parotid gland (PG) and sub-mandibular gland (SMG) cultures. We identified distinctive co-expression of proteins, such as KRT1/5/13/14/15/17/23/76 and 79, exclusively observed in rare, scattered salivary ductal basal cells, indicating the potential de novo source of SGSPC. We also identified an entire class of peroxiredoxin peroxidases, enriched in PG than SMG, and attendant H_2_O_2_-dependent cell proliferation in vitro suggesting a potential role for PRDX-dependent floodgate oxidative signaling in salivary homeostasis. The distinctive clinical resources and research insights presented here offer a foundation for exploring personalized regenerative medicine.

## Introduction

Major salivary glands (SG), namely parotid (PG), sub-mandibular (SMG), and sublingual, play an important role in oral health. These glands have multiple epithelial cell types contributing to dynamic composition of saliva ^1^. The watery secretion of PG, the mucous richness of sublingual, and the versatile watery/mucous mix of SMG highlight their unique contribution to an individual’s quality of life. Salivary dysfunction leading to xerostomia or hyposalivation (persistent dry mouth due to reduced saliva flow) is associated with damages to salivary cells and/or its microenvironment and a blockade of its normal function resulting in a wide range of health consequences.

Salivary dysfunction increases risk of dental caries, gum disease, and oral infections, affects fundamental activities like chewing and swallowing, and influence speech articulation. It can lead to halitosis, oral infections such as oral candidiasis, altered taste perception, and digestive issues. Reduced saliva secretion contribute to oral discomfort, a sensation of stickiness, and irritation, highlighting the importance of addressing salivary dysfunction for the well-being of individuals ^2^.

Salivary dysfunction can result from conditions, including autoimmune disorders like Sjögren’s Syndrome, rheumatoid arthritis, and systemic lupus erythematosus, as well as cancer treatments like head and neck radiation therapy ^3^, PSMA-targeted radionuclide therapy ^4^ and immunotherapies ^5^, and certain medications may cause dry mouth as a side effect. Conditions like diabetes, viral infections (mumps, HIV, hepatitis), aging, trauma, surgery-related nerve damage, dehydration, and salivary gland tumors can also contribute to a decline in salivary function, leading to dry mouth. Despite decades of research in salivary gland function using animal models ^6^, a curative treatment for salivary dysfunction remains elusive ^7^.

Regenerative medicine holds considerable promise in the context of SG dysfunction. Ongoing research explores the use of stem cells (SC) and tissue engineering to regenerate damaged or irradiated salivary glands. Transplantable cells capable of restoring saliva function have been identified in recent years, paving the way for regenerative therapies ^8–10^. These approaches aim to address the consequences of salivary dysfunction, such as xerostomia, by promoting the repair and restoration of functional salivary tissues, but lack of community accessible clinically annotated salivary specimen suitable for SC investigation, which is a major impediment to development of salivary regenerative medicine. To address this gap, we have established a salivary biobank at the Mayo Clinic from patients undergoing head and surgery as well as from autopsies and clinical data and from individuals contributing to our biobank.

The stemness, plasticity and transplantability of salivary cells are not well understood. Studies in exocrine mammary gland with similar epithelial structures suggest that non-transplantable differentiated myoepithelial and luminal cells following short-term in vitro culture or genetic manipulation exhibit stemness, displaying both transplantability and bilneage regeneration ^11–13^. Studies in mouse salivary glands indicate the heightened proliferation potential of lineage-committed differentiated cells, such as acinar cells ^14^, and upon radiation injury, various ductal progenitor cells play a role in maintaining tissue homeostasis ^15^. Therefore, human specimen resources and protocols to investigate these diverse salivary cell types and their regenerative activity and transplantability is essential to develop salivary regenerative medicine.

Herein, we describe an optimal protocol to isolate, expand, and quantitatively investigate salivary cell immunophenotypes and growth properties in 2D and 3D-matrigel. We developed rigorous methods to isolate salivary gland stem and progenitor cell (SGSPC) by flow cytometry using CD24/CD49f/EpCAM+ and PSMA-phenotype. We further generated a unbiased LC-MS/MS-based first known single-cell proteomic map of SGSPC-enriched single cell population. This single-cell proteomic map identified distinct protein co-expression of keratins including KRT1/5/13/14/15/17/23/76 and 79, which are found in rare, scattered salivary gland basal duct cells suggesting their potential role as the tissue origin of SGSPC. Our unique tissue-organoid biobank and novel quantitative approach to study SGSPC provide a robust framework for advancing personalized regenerative medicine research targeting salivary gland dysfunction.

## Results

### Establishment of a Comprehensive Living Salivary Tissue-Organoid Biobank

Using established method from living mammary tissue-organoid biobanking for stem and progenitor cell investigation ^16–30^, we established a clinically annotated salivary biobanking protocol involving salivary tissue collection, mechanical (manual) and gradual enzymatic dissociation, and cryopreservation of intact 2-4 mm salivary units (referred to as ‘salivary tissue-organoids’) (see **Methods**) (**Table 1** and **Figure 1A-J**). For each specimen, the biobank also generates two tissue fragments: one snap-frozen tissue and another for formalin-fixed paraffin-embedded block preparation; and all specimens were tracked using the Mayo Clinic’s Research Laboratory Information Management System (**Figure 1A**). Surgical resections of SMG and PG glands were performed on consenting patients aged 18 and older undergoing head and neck surgery at Mayo Clinic Jacksonville. Tissue samples from deceased individuals were obtained during autopsies at Mayo Clinic Rochester.

**Figure 1.**
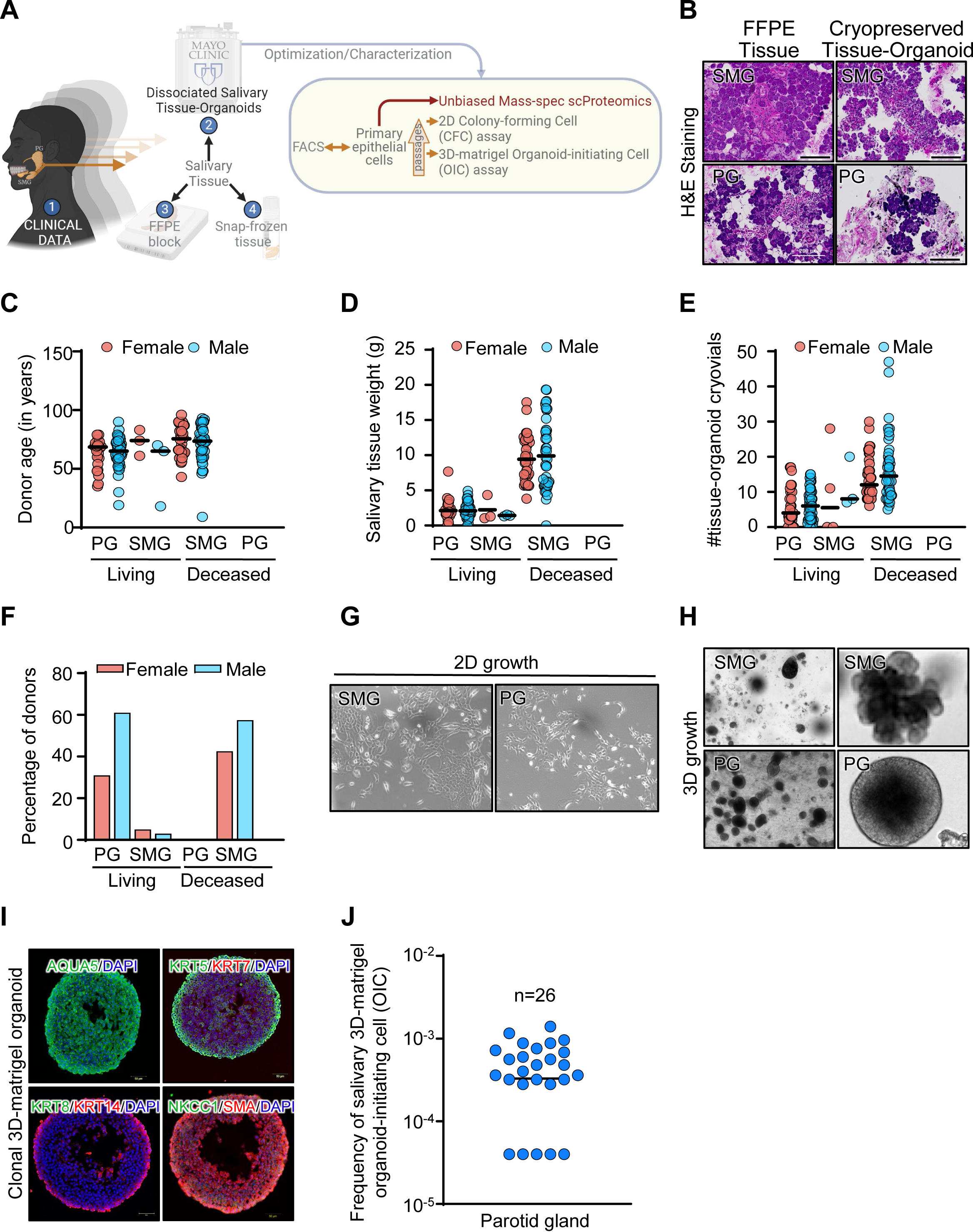
Salivary Tissue-Organoid Biobanking Preserves Primitive Salivary Cells. **A)** Schematic workflow outlining the steps involved in the processing of PG and SMG tissues to generate tissue organoids, as well as their cryopreservation and biobanking from living or deceased male and female donors. **B)** H&E staining of SMG and PG tissues and their corresponding tissue organoids show characteristic ductal and acinar structures (100x magnification). **C)** Plot showing the age distribution among PG and SMG donors, considering both living and deceased individuals of both genders in the biobank. **D)** Plot depicting the weight distribution of PG and SMG tissues received at the laboratory, considering both living and deceased donors of both genders **E)** Plot illustrating the quantity of patient-derived tissue-organoid cryovials obtained from PG and SMG tissue masses received at the laboratory, considering both living and deceased donors of both genders. **F)** Bar plot illustrating the distribution of PG and SMG tissue organoids in the biobank, considering donors from both living and deceased individuals of both genders**. G)** Representative brightfield images of cultured SMG and PG cells isolated from respective cryopreserved tissue-organoids (100x magnification). **H)** Representative brightfield images of SMG and PG organoids cultured in 3D-matrigel (40x and 100x magnification) derived in culture following dissociation of respective tissue organoids. **I)** Representative immunofluorescence staining of 3D-matrigel-derived PG organoids with DAPI, Aquaporin 5, NKCC1, SMA, KRT5, KRT7, KRT8 and KRT14 antibodies. **J)** Plot showing 3D OIC frequency from cryopreserved salivary gland tissue organoids.

**Table 1.**
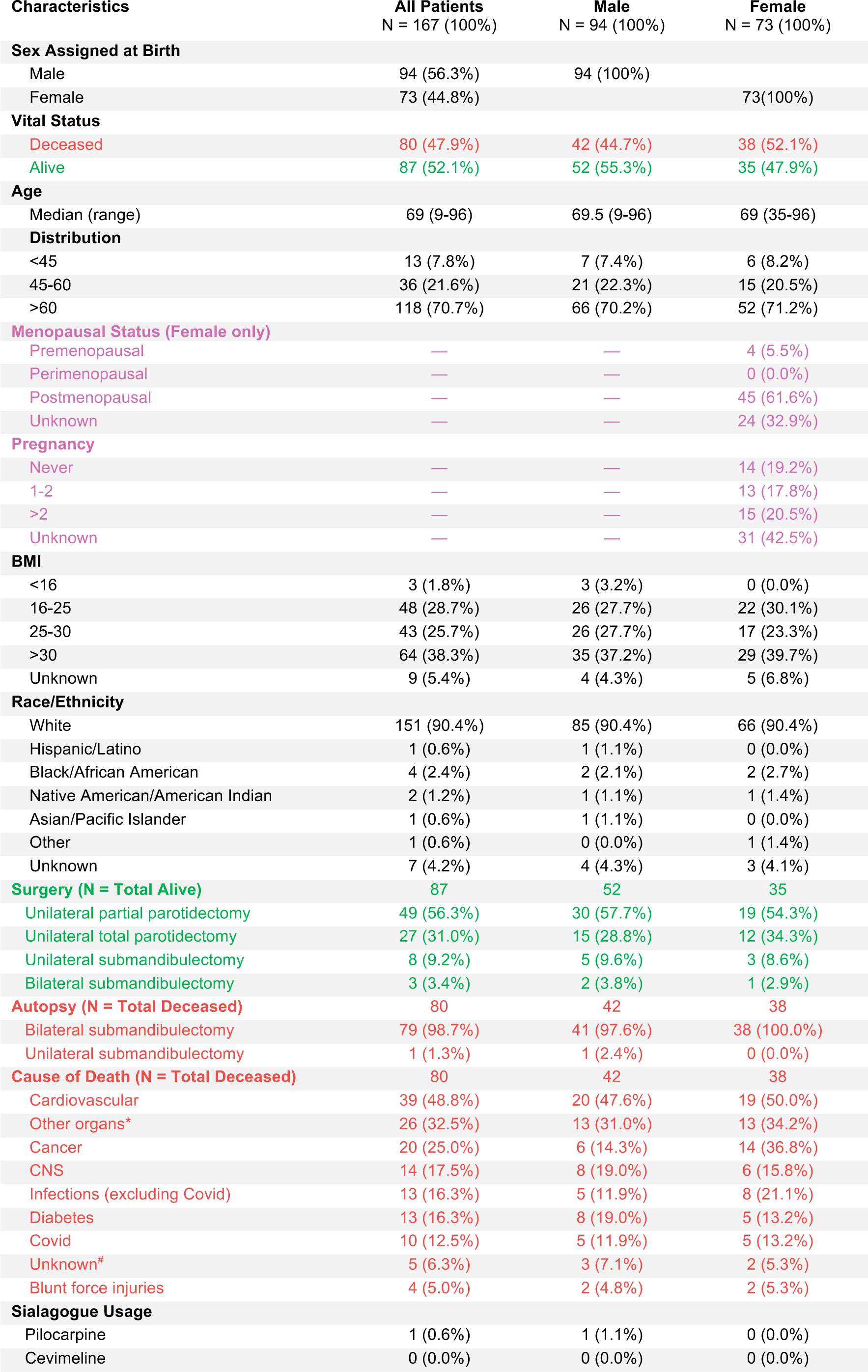
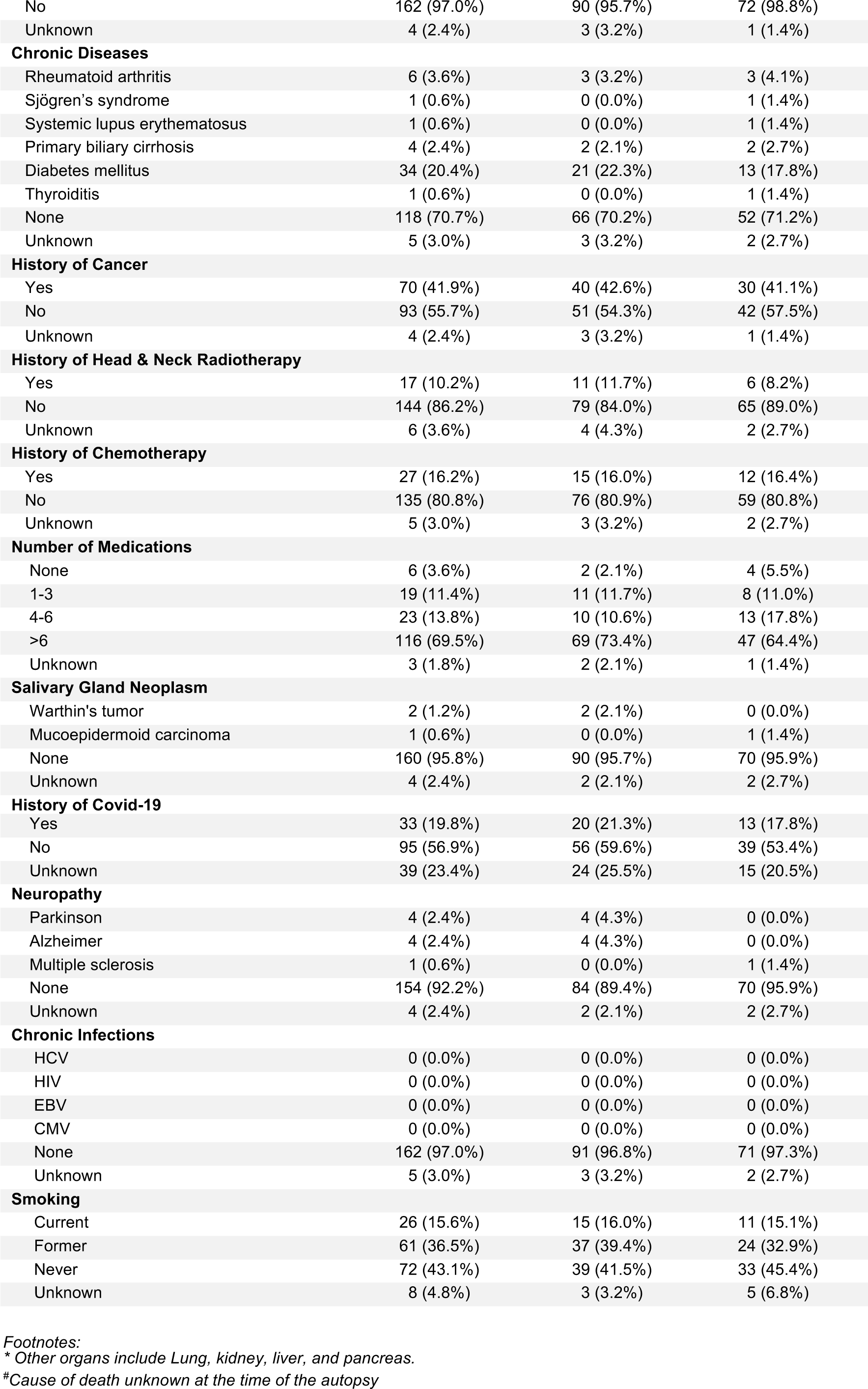
Donor Characteristics in the Mayo Clinic Salivary Regenerative Biobanking.

A total of 249 specimens were collected from 167 donors, with 44.8% obtained from female individuals representing diverse age, race, ethnicity, medical history, various exposures and mortality status, all of which contributed to the biobank (**Table 1**). Cardiovascular conditions were the primary cause of death (48.8%), followed by other organ-related issues (32.5%). Notably, 15.6% were current smokers, and 36.5% were former smokers. Various chronic conditions, such as cancer, head and neck radiotherapy, and chemotherapy treatment, were prevalent among donors. The biobank included samples from patients with salivary gland neoplasms, diabetes mellitus, and rare conditions like Sjögren’s disease. A significant portion had a history of COVID-19 (for details refer to **Table 1**).

The epithelial structures were preserved within cryopreserved tissue-organoids, as confirmed by hematoxylin and eosin staining (**Figure 1B**). Further, the tissue-organoids were dissociated to yield single cells and were evaluated for the presence of primitive cells with proliferative activity using 2D culture and 3D-matrigel organoid culture system that supported clonal development (see **Methods**). In particular, single cell-derived organoids have emerged as systems to approximate progenitor activity in vitro in many tissue types ^31^ including SG ^8^ and represent a valuable tool to evaluate the developmental activity of salivary cells ^32^. A characteristic epithelial morphology and lumenized acinar structures in 2D culture and 3D-matrigel organoid cultures, respectively (**Figure 1G-I** and **Supplementary Figure S1A-B**). From 167 donors, we generated 1,719 tissue-organoid cryovials including 1,031 from male and 688 from female donors. Using the 3D-matrigel organoid-initiating cell (3D OIC) assay system the frequency of SGSPC was estimated to be ∼1 in 1700 isolated single cells from cryopreserved salivary tissue-organoids (**Figure 1J**). This comprehensive biobanking protocol establishes a valuable resource for advancing research in salivary regenerative medicine. For details on accessing biobank specimens and data, kindly refer to the **Methods** section.

### Immunophenotyping Salivary Cells Identifies Rapid Selection of Epithelial Cells In Vitro

We next employed FACS to immunophenotype cells. Hereto, we conducted an in-depth analysis of epithelial biomarker expression in single cells obtained from freshly dissociated PG and SMG tissue-organoids. Additionally, we examined cells from 2D cultured derivatives across various passages in media-1 and media-2. Using cell surface markers reported in the literature and human protein atlas, we developed an antibody cocktail targeting endothelial CD31, hematopoietic CD45, stromal CD34, epithelial markers EpCAM, CD24, and CD49f, and *FOLH1*-encoded prostate specific membrane antigen (PSMA), somatostatin receptor (SSTR)2, and DAPI. We performed FACS, depleting endothelial, hematopoietic, and dead cells, and subsequently gated viable cells for epithelial biomarker expression ^8, 18, 33–37^ (**Figure 2A**).

**Figure 2.**
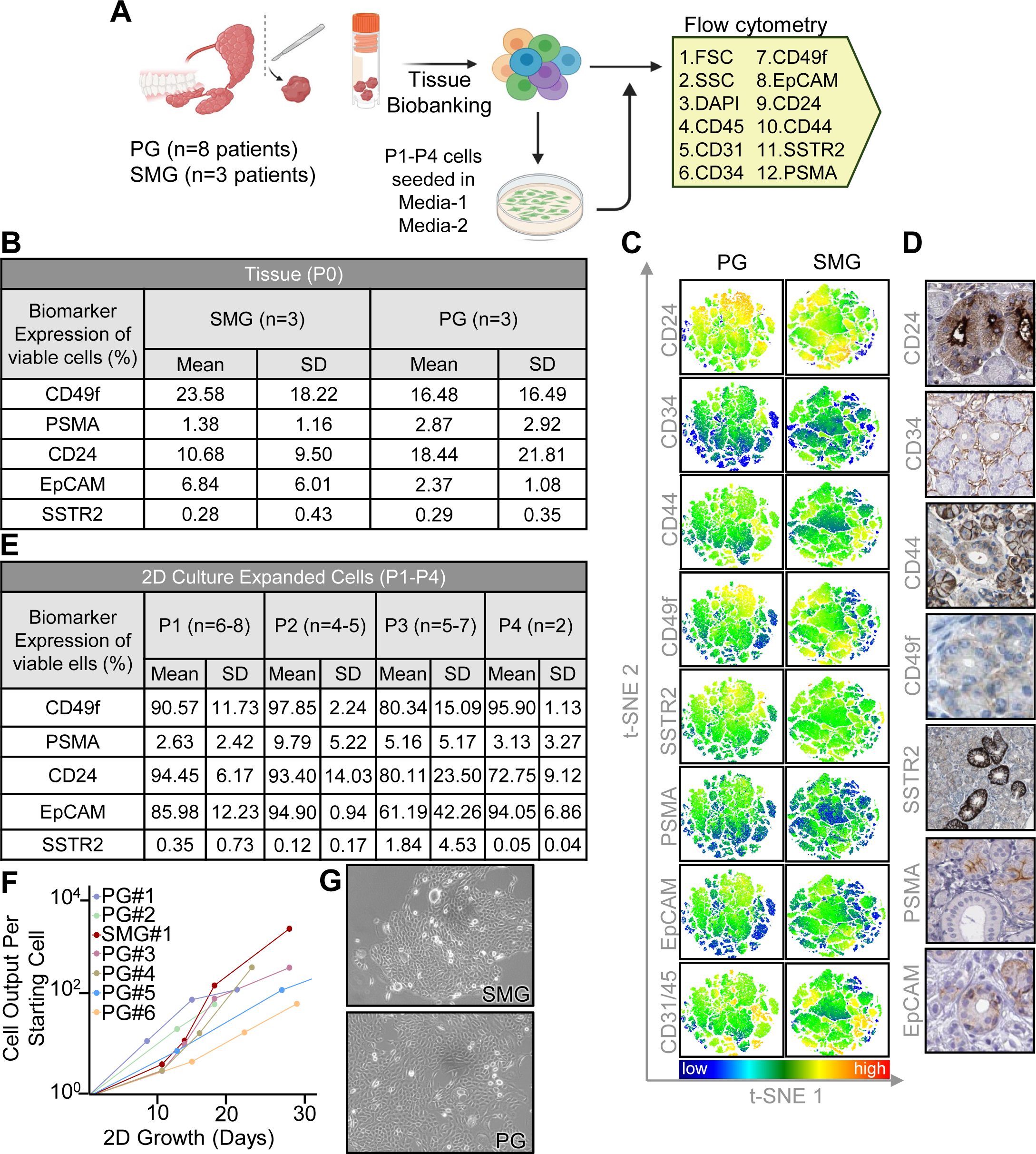
Immunophenotypic Analysis of Salivary Tissue-Organoids. **A)** Experimental design of FACS-based characterization of cryopreserved salivary gland tissue and culture-expanded salivary cells derived from previously cryopreserved tissue. **B)** FACS-based biomarker characterization of unmanipulated salivary gland tissue. **C)** T-distributed stochastic neighbor embedding (t-SNE) plots and histology of PG and SMG tissue by biomarker of interest. Coloring correlates with expression intensity. Red indicates high expression, while blue signifies the absence of the cell marker expression. **D)** Representative immunohistochemical staining showing expression of various biomarkers in salivary glands (Source: Human Protein Atlas). **E)** FACS-based biomarker characterization of 2D-culture expanded cells derived from cryopreserved tissue. **F)** Salivary cells in culture system can be expanded through four passages. **G)** Brightfield image of salivary cell morphology in culture system.

The cell surface epithelial marker expression profiles of freshly dissociated salivary tissue-organoids (CD49f: SMG=23.58%, PG=16.48%; CD24: SMG=10.68%, PG=16.48%; EpCAM: SMG=6.84%, PG=2.37%; PSMA: SMG=1.38%, PG=2.87%, SSTR2: SMG=0.28%, PG=0.29%) were not significantly different suggesting that the dissociation protocol enabled purification of primary salivary epithelial cells with shared immunophenotypes in the two major salivary gland (**Figure 2B**). The expression of these markers remained consistent between genders (**Supplementary Figure S2**).

In 2D culture, salivary cells from SMG and PG exhibited robust expansion for up to four passages. FACS analysis across P1 to P4 passages demonstrated ∼90% cells expressing epithelial markers CD24, CD49f and EpCAM, and lacking expression of PSMA (∼5%) and SSTR2 (<1%) across PG and SMG cultures. Hematopoietic (CD45+) and non-epithelial stromal (CD31/CD34+) cells were found to be poorly detected in cultured cells, indicating rapid selection for epithelial cells under these conditions. Notably, an initial increase in PSMA+ cells was observed in culture, reaching approximately 10% at passage 2 before declining in subsequent passages. SSTR2 expression remained consistently low throughout the culture period, suggesting enrichment of a subset of cells likely to be ductal in origin (**Figure 2C-D** and **Supplementary Figures S3-S4A-C**). In media-1 and media-2, the majority of cultures reached passage 3 by three weeks, while maintaining their characteristic epithelial morphology throughout the entire culture period (**Figure 2E-G**). This comprehensive flow cytometry analysis provides insights into the dynamic expression patterns of salivary epithelial biomarkers in cryopreserved tissue organoids within the biobank and after in vitro expansion, revealing a significant bias toward the selection of ductal epithelial cells.

### Functional Analysis Reveals Distinct Salivary Progenitor Populations In Vitro

We performed a comprehensive assessment of the cryopreserved tissue organoids to test the proliferative epithelial cells in 2D culture enriched for SGSPC. We fractionated salivary cells based on CD24, PSMA, EpCAM, and CD49f markers and subjected bulk cells or distinct phenotypic fractions to optimized 2D colony-forming cell (2D CFC) and 3D-matrigel organoid-initiating cell (3D OIC) assays by seeding cell densities that generate clonal outgrowths (see **Methods**) ^21^. To further understand the growth dynamics, we derived primary salivary cells from cyropreserved salivary tissue-organoids and cultured using media-1 ^38^, both with and without fetal calf serum (FCS), or in N2 media ^9^. Cultures were conducted in dishes with or without collagen-coated surfaces ^17^. Notably, collagen coating had little or no impact on growth in media-1, while cells in N2 media demonstrated sensitivity to collagen coated surfaces (**Supplementary Figure S5A-C**). Despite marginal improvements with the inclusion of FCS, we opted for media-1 with FCS and no collagen coating for this study.

The 2D CFC assays revealed three distinct colony morphologies, each representing unique characteristics—spindle (myoepithelial-like), epithelial (ductal-like), and bridge-forming (squamous-like) colonies (**Figure 3A-B**). The presence of three distinct clonotypes implies three distinct progenitor origins. The persistence of all clonotypes indicates that these progenitors not only consistently cryopreserved in salivary tissue-organoids but also thrive under in vitro culture conditions. Presence of all three 2D CFC types in FACS-purified cells expressing CD24, EpCAM or CD49f and lack of PSMA and SSTR2 indicated that these distinct progenitor types have common immunophenotypes. A prior study, utilizing primary human salivary cells immortalized with origin defective DNA of SV40, has reported similar cellular morphologies in cultures ^39^.

**Figure 3.**
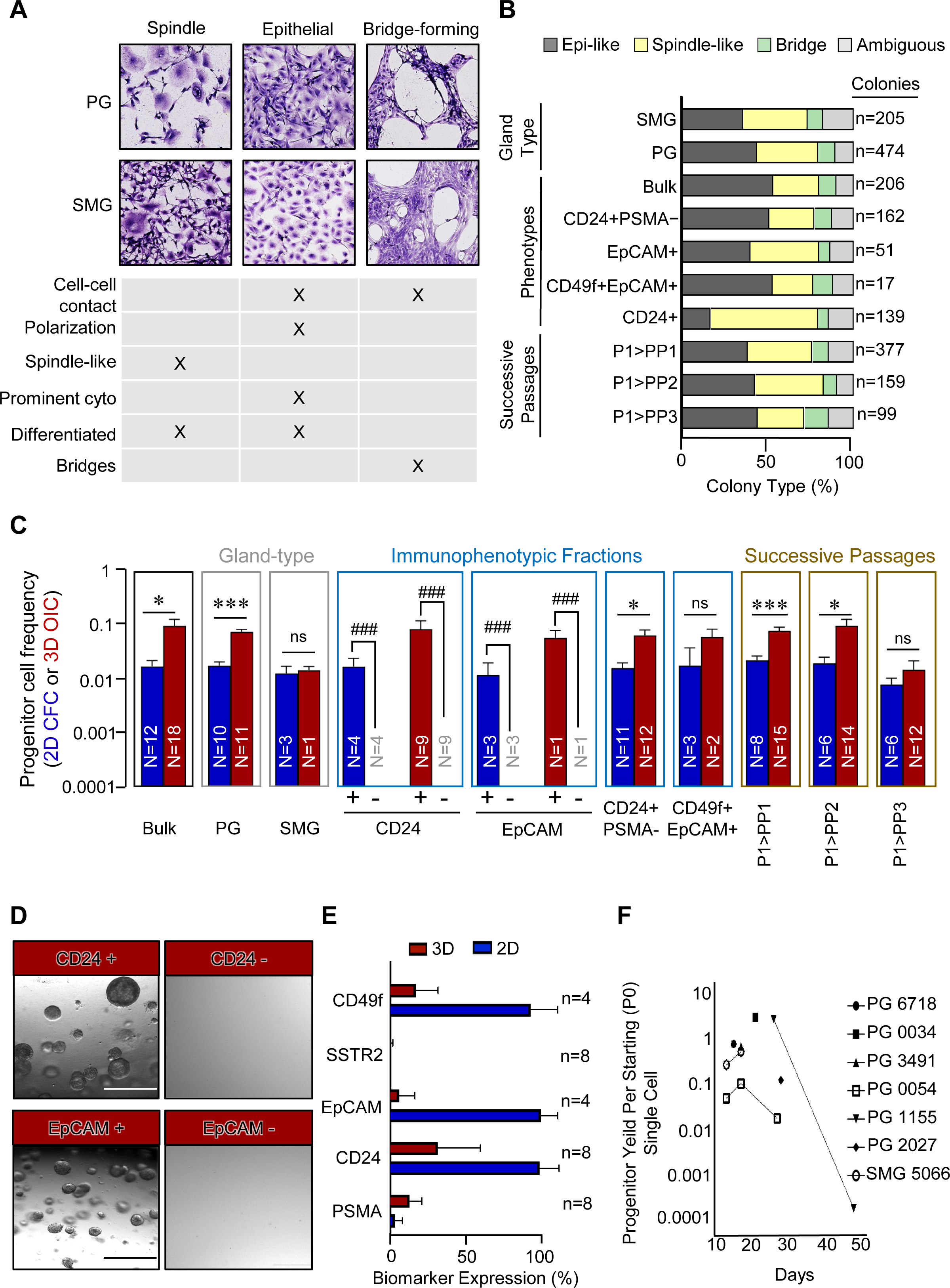
Characterization of Salivary Progenitor Activity in 2D and 3D Assays. **A)** Giemsa staining of salivary colonies obtained in 2D colony-forming cell (CFC) assay showing three distinct morphologies under the microscope. **B)** Plot showing the proportions of each of the three distinct colony types in 2D CFC assays when cells are plated, based on salivary gland type, immunophenotype, or in successive passages. **C)** Plot showing the frequency of CFC and 3D-matrigel OIC when plated based on salivary gland type, immunophenotype, or in successive passages in the 2D CFC and 3D-matrigel organoid assays. **D)** Brightfield microscopy images showing 3D-matrigel organoids obtained from FACS-purified CD24+ and EpCAM+ salivary cells, but not from CD24- and EpCAM-salivary cells. **E)** Immunophenotypic characterization of single cells obtained from 3D-matrigel organoids and 2D culture expanded cells from human patient salivary gland tissue organoids. **F)** Plot showing progenitor yield per starting single PG and SMG cell in culture over time. (*, **, and *** indicate p-values < 0.05, 0.005, and 0.0005, respectively. #, ##, and ### represent p-values < 5 x 10^-7^, 5 x 10^-10^, and 5 x 10^-13^, respectively).

Furthermore, our investigation revealed detection of upto 18% of 3D-matrigel organoid-initiating cell (OIC) population for PG cells at passage 3. The average OIC was 3.3% across diverse tissue sources and in vitro culture passages (**Figure 3C** and **Supplementary Figure S4C**). Notably, the progenitor frequencies observed in the 3D-matrigel OIC assay were consistently higher compared to the 2D colony-forming cell (CFC) assay. Similarly, we detected a maximum of 5% in 2D CFC population with in EpCAM+ fraction of PG cells and on average 1.6% across diverse tissue sources and in vitro culture passages (**Figure 3C** and **Supplementary Figure S4C**).

Unlike the 2D CFC assay, we did not observe visible morphological differences in the 3D-matrigel outgrowths (**Figure 3D** and **Supplementary Figure S6**). Additionally, the significant association among salivary gland stem/progenitor cell (SGSPC) markers, including CD24, EpCAM, and CD49f (p-value < 0.0001), was evident (**Supplementary Figure S7**). In-depth examination of cells derived from 3D-matrigel organoids, as opposed to those expanded in 2D culture, revealed distinct expression patterns of biomarkers, underscoring the intricate and dynamic nature of the differentiation process within the 3D-matrigel environment (**Figure 3E**). Additionally, it is noteworthy that the progenitor yield per starting single PG and SMG cell remained comparable over time in culture (**Figure 3F**). Together, this implies that the methodologies outlined in this study consistently generate salivary gland stem/progenitor cell numbers, emphasizing the robust selection of the proliferative CD24/EpCAM/CD49f+ SGSPC subset by the specific culture environment.

### Induction of PSMA Expression during Salivary Progenitor Cell Differentiation

The significance of PSMA is underscored by the emergence of PSMA-targeted cancer therapies, which are known to cause salivary dysfunction. Therefore, there is a topical interest in understanding the role of PSMA during salivary development. First, we FACS-purified both PSMA+ and PSMA-cells from bulk cultures and assessed their growth properties in 2D CFC and 3D OIC assays (**Figure 4A-C** and **Supplementary Figure S6**). Notably, a maximum of 2.75% (average 1.15%) and 22% (average 3.74%) of PSMA-cells exhibited robust 2D CFC and 3D OIC activities respectively, while PSMA+ cells were devoid of any growth potential. The PSMA expression is restricted to acinar secretory cells in situ (**Figure 2D**). Remarkably, PSMA expression was exclusively localized to the apical membrane of secretory cells within terminal acinar structures, conspicuously absent from ductal regions (**Figure 2D**). Together, these findings imply a plausible hierarchical differentiation model, proposing that PSMA-SGSPC may serve as progenitors for non-cycling (differentiated) PSMA-positive cells with an acinar fate (**Figure 4D**). To investigate this hypothesis, PSMA-cells were FACS-purified and cultured in 2D or 3D conditions (**Figure 4E-G**). Subsequent FACS analysis revealed a gain in PSMA expression in both 2D culture-expanded and 3D-matrigel organoid-derived cells, supporting the notion of PSMA-cells transitioning to a PSMA+ phenotype. The induction of PSMA expression in cells generated in 2D and 3D-matrigel cultures was observed in all patient samples tested (**Supplementary Figure S8A-C**) suggesting spontaneous but poor differentiation in these culture environments.

**Figure 4.**
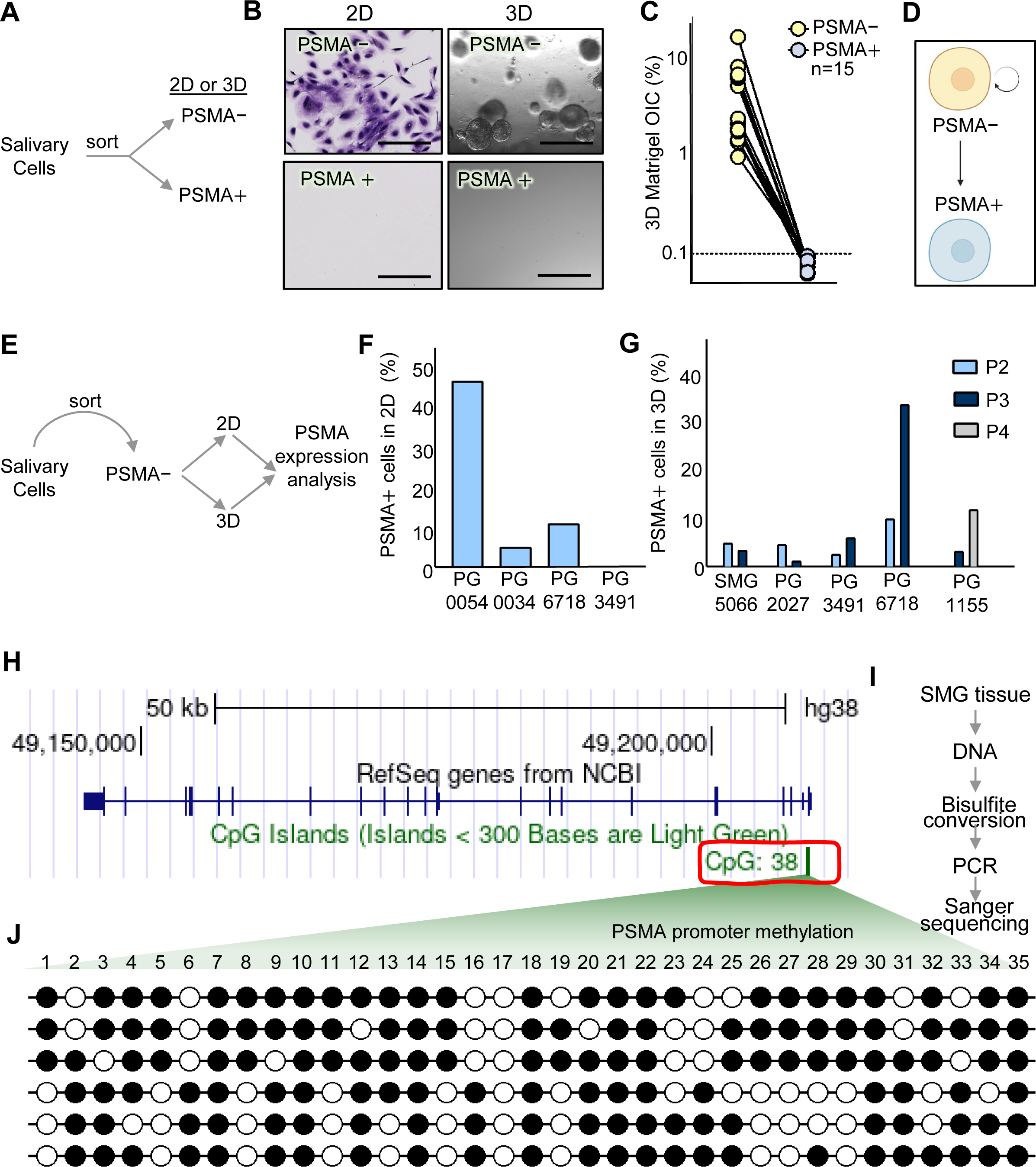
PSMA is a SGSPC Differentiation Marker. **A)** Design of 2D CFC and 3D-matrigel organoid Matrigel experiments to assess clonogenicity of PSMA- and PSMA+ cells. **B)** PSMA-cells demonstrate clonogenic capacity in 2D CFC and 3D-matrigel organoid assays, while PSMA+ cells do not. **C)** PSMA-cells purified from 15 salivary gland patient samples, unlike PSMA+ purified cells, generate organoids in 3D culture. **D)** Proposed hierarchical lineage model by which PSMA+ cells differentiate from the more primitive, stem/progenitor-like, self-renewing PSMA-cells. **E)** Experimental design of 2D and 3D FACS-based characterization of growing PSMA-salivary cells to gauge PSMA+ cell generation. **F)** PSMA-cells plated in 2D culture generate PSMA+ cells at confluence. **G)** 3D-matrigel organoids derived from PSMA-cells give rise to PSMA+ cells through multiple passages. **H)** Illustration of CpG islands in human PSMA promoter region. **I)** Schematic workflow of bisulfite conversion of 35 of 38 CpG, PCR and sanger sequencing of DNA obtained from SMG. **J)** Hypermethylation (∼70%) of CpG in *PSMA* promoter in SMG tissue. Each circle indicates individual CpG dinucleotides. White and dark circles represent unmethylated and methylated CpG, respectively.

Gene promoter DNA methylation during cellular differentiation, involving the addition of methyl groups to cytosine nucleotides adjacent to guanines (CpG dinucleotides), is a widespread epigenomic mechanism of gene regulation. A careful examination of the PSMA promoter revealed that it contains a CpG island (**Figure 4H**). Thus, we asked whether PSMA is targeted for methylation-mediated silencing in human salivary tissue. We devised a methylation-specific PCR assay to examine PSMA promoter methylation, followed by bisulfite sequencing to validate the hypermethylation of CpG islands within the PSMA promoter in DNA from snap-frozen SMG tissue (**Figure 4I**). Our experiments validated PSMA promoter hypermethylation in salivary tissue. Our investigation reveals a compelling association between PSMA-primitive salivary cells and the emergence of differentiated PSMA+ salivary acinar secretory cells, suggesting a potential regulatory role for DNA methylation during acinar differentiation.

### Unbiased Mass Spectrometry-Based Single-Cell Proteomic Profiling of Primitive Salivary Cells

To gain molecular insights into SGSPC, we performed proteome profiling of single cells using trapped ion mobility spectrometry (TIMS) in conjunction with a time-of-flight mass spectrometer operated in parallel accumulation-serial fragmentation (PASEF) mode ^40, 41^. This cutting-edge methodology enhances proteome coverage from low-input samples, enabling the generation of label-free, unbiased liquid Chromatography with tandem mass spectrometry (LC-MS/MS)-based single-cell proteomic profiles of SGSPC-enriched single cells (**Figure 5A**) ^42, 43^. To accomplish this, we used our strategy to enrich for progenitors FACS sorting viable (DAPI-) CD45/CD31-EpCAM+ progenitor-enriched cells (referred hereafter as progenitor-enriched single cells) through stringent gating procedures. Subsequently, these single cells were isolated into a 384-well plate and underwent peptide digestion. These peptides analyzed using a data-independent acquisition method using PASEF (diaPASEF) with a 25 m/z isolation window. The approach consistently facilitated the detection of ∼675 proteins per progenitor-enriched 71 single cells in PG and SMG, respectively (**Figure 5B** and **Table S1**). In fact, our analysis identified proteins that were highly enriched and shared between both SMG and PG progenitor-enriched single cells (**Figure 5C**). Through further analysis of the proteomics, we also identified protein signatures that were unique to SMG (**Figure 5D**) and PG (**Figure 5E**) cells. Therefore, our findings substantiate the existence of shared protein expression patterns among SGSPC, underscoring a common functional profile for these primitive salivary cells irrespective of origin.

**Figure 5.**
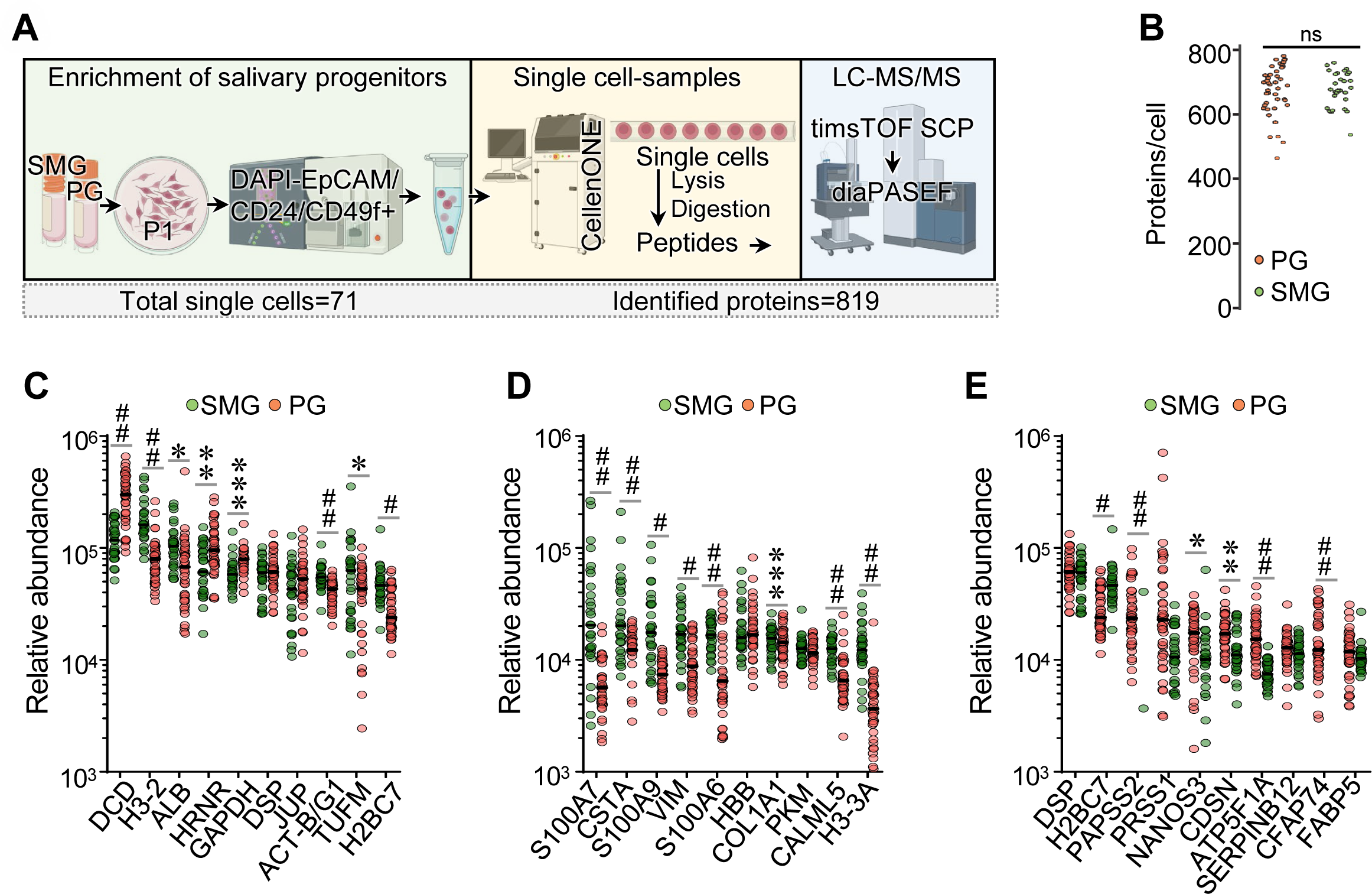
Unbiased LC-MS/MS-Based Single-Cell Proteomic Profiling of Purified Primitive Salivary Cells. **A)** Overall workflow for single cell proteomics. Single cells (P1) from PG and SMG cells were isolated by FACS using DAPI-CD45-CD31-EpCAM+ immunophenotype, lysed and digested in 384-well plate using cellenONE platform. Mass spectrometry data of each single cell were acquired in diaPASEF mode using the timsTOF SCP mass spectrometer. A total of 819 proteins across 71 single cells were identified. **B)** Number of identified proteins per cell from SMG and PG P1 cultures. On average, 674 proteins and 678 proteins per single cell were identified from PG and SMG, respectively. **C)** A plot showing protein counts per cell, highlighting the 10 highly expressed and shared proteins within the progenitor-rich single cells of both SMG and PG. **D)** A plot showing protein counts per cell of 10 most abundantly detected proteins in progenitor rich cells of SMG relative to PG. **E)** Plot showing protein counts per cell of 10 most abundantly detected proteins in progenitor rich cells of PG relative to SMG (*, ** and *** indicate p-values < 0.05, 0.005 and 0.0005, respectively. #, ## and ### represent p-values < 5 x 10^-7^, 5 x 10^-10^ and 5 x 10^-13^, respectively).

### Single-Cell Proteomics Identifies Putative Origin of Primitive Salivary Cells

Cytokeratins are structural proteins specific to epithelial cells, and their expression patterns can be used to identify and characterize different cell types within epithelial tissues ^44^. Therefore, we analyzed PG and SMG single-cell proteomics data to identify cytokeratins distinctly associated with SGSPC. Our comprehensive analysis identified a total of 35 cytokeratins expressed in SMG and PG cells. However, 7 cytokeratins (KRT1, KRT2, KRT10, KRT77, KRT18, KRT7, and KRT3) were significantly upregulated in SMG cells compared to PG cells, while 9-10 cytokeratins (KRT8, KRT84, KRT71/73, KRT31, KRT33B, KRT36, KRT14, KRT27, and KRT37) were significantly upregulated in PG cells compared to SMG cells (**Figure 6A**). We validated expression of these cytokeratins in salivary glands using the human protein atlas and many of these cytokeratins were localized in ‘scattered’ basal cells within salivary ducts (**Figure 6B-C**) with some overlap with myoepithelial cells within salivary acini (**Supplementary Figure S9**) suggesting their potential developmental linkage.

**Figure 6.**
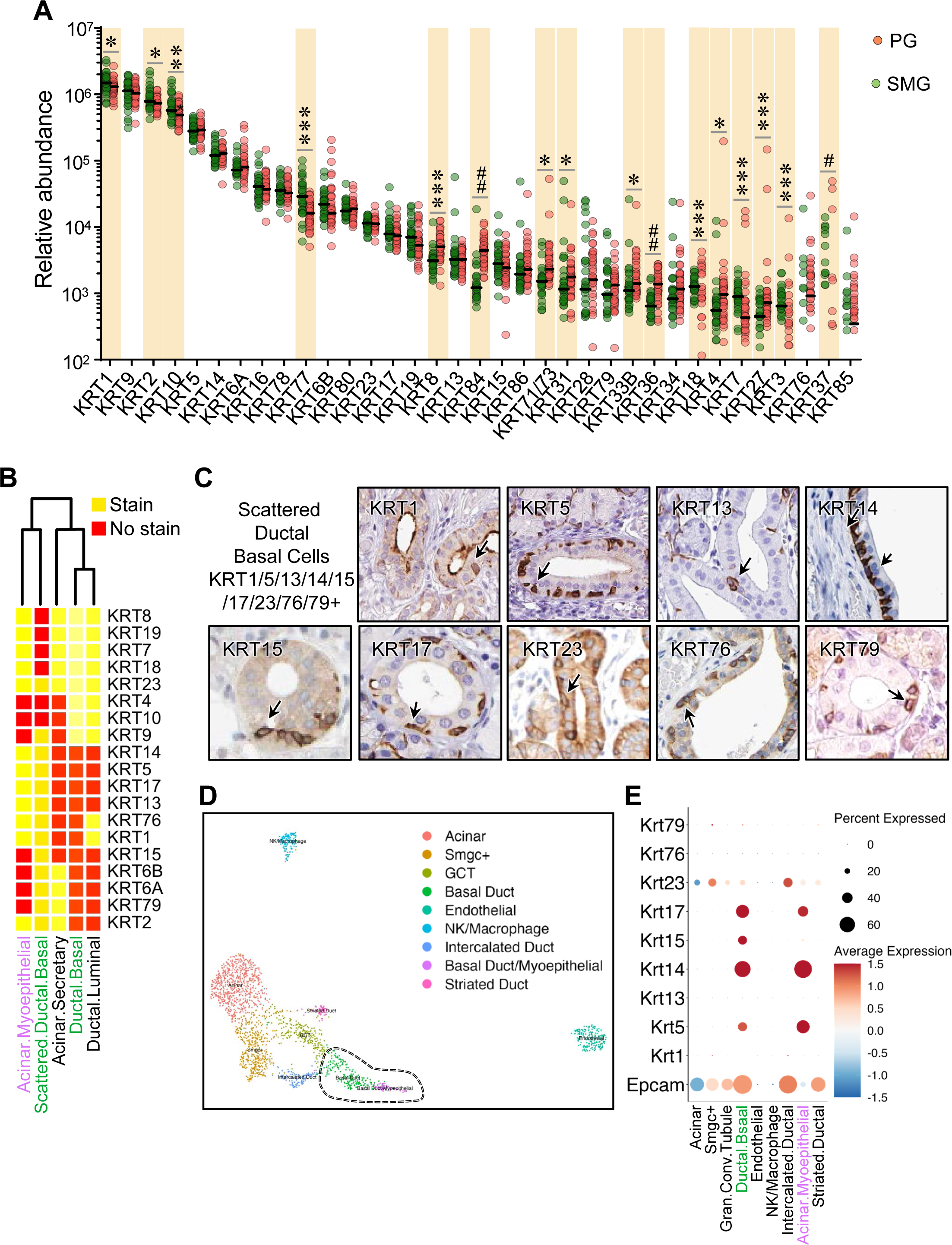
Unbiased LC-MS/MS Identifies Salivary Stem/Progenitor-specific Cytokeratin Profile. **A)** A plot showing the spectral counts of keratin proteins detected per individual cells (P1) from PG and SMG cells were isolated by FACS using DAPI-CD45-CD31-EpCAM+ immunophenotype enriching for SGSPC activity. **B)** The relationship between the keratin expression profile identified through single-cell proteomics in SGSPC and the salivary epithelial cell-type specific expression patterns in the immunohistochemistry-based map of human protein expression profiles explored through the Human Protein Atlas (https://www.proteinatlas.org/). **C)** SGSPC-expressed keratins map to scattered ductal basal cells in salivary glands (https://www.proteinatlas.org/). **D)** UMAP-based cluster analysis of mouse SMG scRNA-seq data showing salivary cell clusters. **E)** Dot plot showing the expression of murine analogs of human SGSPC-expressed cytokeratins (and EpCAM) across mouse salivary cell types (*, ** and *** indicate p-values < 0.05, 0.005 and 0.0005, respectively. #, ## and ### represent p-values < 5 x 10^-7^, 5 x 10^-10^ and 5 x 10^-13^, respectively).

We next searched for murine analogs of these human cytokeratins in publicly available single-cell transcriptome databases and consistently found their association with the ductal-basal cell compartment of salivary cells with multiple cytokeratins overlapping with basal/myoepithelial cell clusters providing supplementary evidence supporting their probable development proximity (**Figure 6D-E**) ^45^.

Our findings reveal a remarkably conserved cytokeratin profile in salivary glands across mammalian species, enabling the identification of rare primitive cells. Utilizing comprehensive keratin profiling via single cell proteomics and scRNA-seq, we present compelling evidence indicating intricate developmental interconnections among ductal basal and acinar myoepithelial cells, potentially exhibiting niche-specific functions ^46^. Moreover, our data suggests that a rare subset of scattered basal cells within ducts, similar to mammary gland, may harbor more primitive functionalities, accessible through our in vitro culture conditions, which are vital for future studies ^47^.

### Primitive Salivary Cells Display Oxidative Signaling

To identify differentially expressed proteins, we first performed principal component analysis (PCA) on the proteins detected in SMG and PG cells. This analysis successfully separated SMG and PG cells into distinct clusters (**Figure 7A**), suggesting inherent biological differences between these cells. Further analysis revealed significant differences in the expressed proteins and identified 94 and 98 differential protein signature (fold-change > 2 and adjusted p-value < 0.01) that defines SMG and PG cells (**Figure 7B-C** and **Supplementary Table S1-S2**). Cellular peroxiredoxins (PRDX) are important components of the cellular defense system against oxidative stress, contributing to the maintenance of cellular redox balance and the prevention of oxidative damage ^48^. Dysregulation of peroxiredoxins or increased oxidative stress in the SG can potentially contribute to conditions such as inflammation, dry mouth (xerostomia), or other oral health issues ^49^. We utilized single-cell proteomics to identify components of oxidative stress pathway in SGSPC. Further analysis revealed several components of oxidative stress pathway including PRDX2 PRDX4 PRDX6 and GLRX to be significantly enriched in PG cells compared to SMG cells (**Figure 7D** and **Supplementary Figure S9**). To functionally validate the role of antioxidant pathways, we treated cultured PG cells with hydrogen peroxide (H_2_O_2_) at various concentrations and evaluated their viability by MTT assay. In fact, the PG cells exhibited resistance to H_2_O_2_ at concentrations as high as 50μM (**Figure 7E**). Taken together, these observations suggest that H_2_O_2_ may influence the oxidation of cellular signaling proteins through peroxiredoxins.

**Figure 7.**
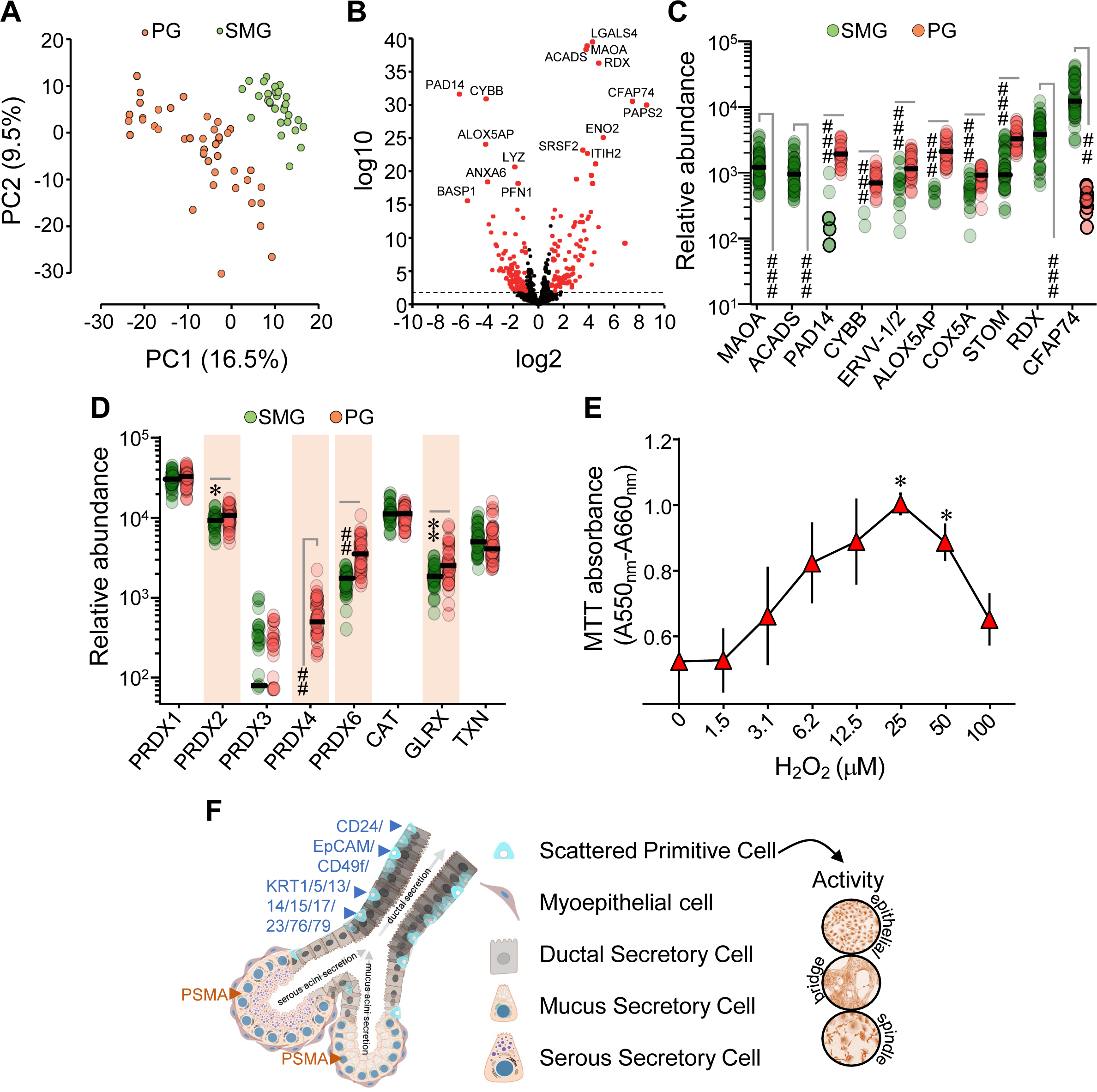
Single-Cell Proteomic Profiling Reveals Floodgate Oxidative Signaling in Salivary Cells. **A)** Principal component analysis of 819 proteins from SMG and PG cells. Each circle represents data from an individual single cell. **B)** A volcano plot showing fold-changes and adjusted p-value of proteins. Differentially expressed proteins (fold-change >2 and adjusted p-value <0.01) are marked in red. **C)** A plot showing top 10 differential proteins between SMG and PG salivary progenitor-rich cells. **D)** Antioxidant enzyme counts per individual cell, illustrating predominantly glutathione-independent peroxiredoxin system in salivary cells. **E)** Cellular metabolic activity in proliferative PG (P1) cells was assessed using the MTT assay, measuring absorbance changes to evaluate cell viability after treatment with varying concentrations of H_2_O_2_ over a 72-hour duration.

## Discussion

We have developed a researched protocol for the first clinically annotated salivary gland biobank at the Mayo Clinic to address the current gap in accessible resources for regenerative medicine research in the United States. Notably, our biobank includes PG and SMG samples from both living and deceased donors, providing diversity in terms of age, race, gender, and disease conditions. Our biobank encompasses specimen from various chronic salivary conditions, including Sjögren’s disease, and neoplasms such as mucoepidermoid carcinoma ^50, 51^ and Warthin’s tumor ^52^. Significantly, our collection includes salivary specimen from patients with history of COVID-19 infection, that may model and understand of SARS-CoV-2 viral replication in the salivary glands and its potential role in oral transmission and COVID-19-associated xerostomia ^32, 53, 54^.

Despite advancements in the understanding of SG development and maintenance, effectively managing SG hypofunction and dysfunction remains a significant clinical challenge. Current treatments are primarily palliative and involve the use of moisturizers such as artificial saliva or saliva substitutes, as well as systemic medications (e.g., pilocarpine or cevimeline) to stimulate salivary secretion ^55^ However, these synthetic agents often fall short in adequately compensating for the natural reduction in saliva production and alleviating associated symptoms, particularly in cases of severe SG damage^56^. Various therapeutic approaches are currently under assessment to address salivary flow restoration in xerostomia patients, with gene therapy being one of them. Ongoing clinical trials are exploring the use of AQP1 gene therapy, employing adeno-associated viruses (AAVs) known for their lower immunogenicity and longer-lasting gene expression compared to adenoviruses ^57, 58^. However, it is worth noting that tissue damaged by radiation therapy may pose greater challenges for gene therapy, as the fundamental cause of SG dysfunction involves extensive disruption or destruction of the gland at both cellular and extracellular matrix (ECM) levels^59^.

Recent clinical observations indicate that irradiating areas containing the highest concentration of SG progenitor cells results in the most substantial reduction in saliva production^60^. These findings suggest that the potential therapeutic approach of SC replacement holds promise for addressing IR-induced xerostomia. One promising avenue of research entails the cultivation of progenitor (or stem) cells *in vitro* to facilitate the production of SG structures, such as salispheres or organoids. These engineered constructs can then be transplanted or infused into compromised tissue to initiate the development of a new functional gland within its native site ^61^. However, despite a substantial body of literature investigating the feasibility of various cell populations for the restoration of SG function, the quest for an abundant and readily accessible source of SG progenitor or stem cells remains ongoing^9, 15, 61, 62^. Nevertheless, a noteworthy development is the first-in-human clinical trial (clinicaltrials.gov identifier: NCT04593589) on autologous SG stem-cell transplantation started in May 2022 undertaken by the University Medical Center Groningen in The Netherlands adding momentum for this research.

The biobank exhibits enrichment for epithelial cells while preserving the heterogeneity of the parental tissue, facilitating the isolation and characterization of transplantable cells with stem/progenitor activity crucial for salivary gland homeostasis. Ongoing studies are exploring these aspects. Viable bulk single cells, purified from cryopreserved specimens, consistently generated organoid structures in matrigel cultures, indicating the presence of cells with primitive activity in our biobanked samples. The lineage hierarchy within SG tissues remains incompletely understood, posing a challenge for the translation and regenerative application of SGSPC ^61^. Our identification of markers, including CD24/EpCAM/CD49f as primitive and PSMA as a differentiation marker, contributes to addressing this gap and may aid future investigations (**Figure 7F**). Notably, recent reports have identified CD49f as a reliable strategy for human SGSPC enrichment by FACS [^63^; validated in our study (**Figure 2C-D**)], and the biobank is poised to support such inquiries in the future.

In this study, we generated the first proteomic map at single-cell resolution of SGSPC-enriched cell population. Our analysis revealed common and unique proteins in SMG and PG cells, including KRT5 and KRT14 associated with salivary progenitor cells ^64, 65^. These keratins serve as essential markers for identifying and isolating stem/progenitor cells in the salivary gland. Notably, KRT5 and KRT19 are recognized progenitor markers in developing salivary glands ^66^. KRT14+ progenitor cells are crucial for maintaining ductal homeostasis in submandibular glands ^15, 67^. Our findings also include KRT7, marking differentiated luminal ductal cells, and KRT76, associated with differentiating layers in stratified epithelia. Understanding these keratins’ roles in various tissues indicates their diverse functions, influencing progenitor cells, ductal homeostasis, and differentiation within salivary glands. Similarly, PRDX were among commonly highly upregulated proteins in SGSPC of SMG and PG except for PRDX4 which was almost exclusively expressed in PG suggesting role for these proteins in regulating floodgate oxidative signaling in salivary glands. Our salivary tissue organoids provide a valuable resource for further exploration of these roles.

In summary, the Mayo Clinic’s salivary tissue-organoid biobanking and research approach to investigate SGSPC, combined with their proteomic maps, represent a novel resource for salivary regeneration research.

## Methods

### Donor eligibility, informed consent process, specimen collection sites and access

The Salivary tissue organoid biobank is an opt-in biobank for which written informed consent is obtained from the participants. The informed consent is reviewed and approved by the Mayo Institutional Review Board to ensure that the consenting process meets all legal requirements. Biobank participants agree to permit the use of samples and/or data from multiple studies, provide access to questionnaires and data from the medical record (including past, present and future details). Moreover, for the Mayo Clinic Salivary tissue organoid biobank, participants allow access to stored clinical samples, and permit the sharing of deidentified data with other researchers. Participants are not contacted for additional studies. The consent document discusses privacy protections and risks involved in participation in the study. Further, the consent discusses the potential for receiving results from projects that use the salivary tissue organoid biobank as a resource. It also provides 2 check boxes (included at the suggestion of the community members), allowing the participant the options of (1) not allowing access to stored clinical specimens for research and (2) not allowing family members access to samples after their death.

Patients at Mayo Clinic who are 18 years or older, who can communicate in English or use interpreters if needed, able to consent, are eligible to enroll in the Salivary Tissue Organoid Biobank. Study materials are provided in English only. Recruitment to the Biobank is primarily conducted in person for prospective participants scheduled for a pre-surgical appointment at Department of Otorhinolaryngology, Mayo Clinic Jacksonville. The enrollment started at Mayo Clinic Jacksonville in September 2020, with active ongoing collections. All requests for specimens and data will be thoroughly reviewed in accordance with institutional policy. For further details, please email corresponding authors.

### Viable salivary specimen processing, quality control and derivatives

Tumor-free clinical residual SG specimens from living donors are obtained from the anatomic pathology laboratory at Mayo Clinic Jacksonville, and SMG tissue from deceased donors is obtained from the autopsy laboratory at Mayo Clinic Rochester. The specimen from living donor is collected in a buffered media containing DMEM (Corning), 5% FCS (Gibco), and 1x penicillin-streptomycin (Gibco) and transported overnight on ice (FedEx) to the Stem and Cancer Biology Laboratory at Mayo Clinic Rochester for further processing and biobanking. Specimens are processed using the mechanical and enzymatic dissociation method. Briefly, tissues were finely minced using sterile blades and dissociated overnight in DMEM F-12 (Ham’s, Gibco) supplemented with 2% wt/vol BSA (Fraction V; Life Technologies), 300 U/mL collagenase (Sigma-Aldrich), and 100 U/mL hyaluronidase (Sigma-Aldrich) at 37°C and 5% CO_2_. Further, epithelial-rich tissue organoids were obtained following centrifugation at 100 × g for 5 min. Subsequently, the tissue organoids were cryopreserved in freezing media containing 50% DMEM (Corning), 44% FCS (Gibco), and 6% DMSO (Invitrogen). Each tissue organoid cryovial is barcoded with sample attributes and are assigned storage using RLIMS (Research Laboratory Information Management System). RLIMS is a web-based application utilized by Mayo Clinic research laboratories to create, annotate, and track samples. Moreover, RLIMS can seamlessly integrate with laboratory sample processing, storing, and delivering structured data on samples. Further, for each specimen, a fraction of tissue-organoids is dropped into DMEM and tested for microbial contamination for 7 days at 37°C and 5% CO_2_ and results recorded.

### Microscopy

Organoids were then stained as previously described ^6^ with primary and secondary antibodies described in **Supplementary Table S3**. Nuclei were counterstained with DAPI Fluoromount-G® (Southern Biotech, Birmingham, AL, USA). All stained slides were imaged with Cytation5 (BioTek Instruments Agilent Technologies, USA).

### Primary salivary cell isolation and expansion

The cryopreserved tissues were partially thawed in a water bath maintained at 37°C and retrieved by centrifugation at 300g for 5 minutes in HBSS buffer supplemented with 2% FCS and 1x penicillin-streptomycin. Single-cell suspensions were then obtained from tissue organoids by serially dissociating with trypsin-EDTA (0.25%, STEMCELL Technologies), 5 mg/mL dispase (STEMCELL Technologies), supplemented with 100 µg/mL DNase1 (Sigma-Aldrich). The cell suspension was passed through a 40-µm strainer and cells were collected by centrifugation ^17^. Viable cells were counted following trypan blue (0.4%, Gibco) staining in Neuber’s chamber and subsequently, cells were 2D cultured in media-1 as described elsewhere ^21^.

### 3D-matrigel salivary organoid-initiating cell (OIC) assay and organoid analysis

Freshly dissociated salivary cells from cryopreserved organoids were evaluated for OIC activity in a 3D-matrigel culture as described previously for human mammary cells ^21^. In brief, a suspension of 25,000 freshly dissociated cells or culture-expanded/FACS-sorted cells ranging from 625 to 5000 cells were prepared in growth factor reduced Matrigel (R&D Systems) and plated in 96-well plates as domes. The domes were allowed to solidify for 15-30 minutes. Subsequently, 200 μL of pre-warmed media-1 and media-2 was added. media-1 is prepared using DMEM/F12 (Ham, Gibco), supplemented with 5% FCS, 10 ng/ml EGF, 10 ng/ml cholera toxin (Sigma), 1µg/ml insulin (Sigma) and 0.5 µg/ml hydrocortisone (Sigma) ^21^. Limited studies were carried out using media-2 but was prepared using Advanced DMEM/F12 (Gibco) supplemented with 1% B27, 10 ng/mL EGF, and SB431542 as published elsewhere ^68^.

### 2D colony-forming cell (CFC) assay

The culture-expanded or FACS purified salivary cells with distinct phenotypes were seeded at densities of 200, 600 and 1800 in a 6-well plate and cultured for 10 days in media-1. The media was removed and cells were rinsed with PBS, fixed using acetone:methanol (1:3), and then stained with Giemsa stain (Sigma-Aldrich) for colony visualization. Colony morphologies were visualized using microscope and classified into various types as described elsewhere^19, 20, 39^.

### Flow cytometry

Salivary cells from cryopreserved tissue-organoids, cultured 3D-matrigel organoids or 2D culture expanded cells at different passages were analyzed by FACS. In order to isolate cells from organoids, the organoid-matrigel pellet was resuspended in 100 μL of Cell Recovery Solution (Corning) and incubated on ice for 30 minutes. The solution was centrifuged at 2500 x g for 10 minutes and the supernatant removed to isolate salivary cells from matrigel. 500 μL of 0.05% Trypsin was added and the solution was mixed by pipetting up and down. 1 mL of HBSS (Gibco) supplemented with 2% FBS was then added prior to a 5-minute incubation at 37°C and subsequent centrifugation to yield single cells. The cells were stained with anti-human fluorochrome-conjugated antibodies listed in **Supplementary Table S4**, and DAPI to identify dead (DAPI+) cells. Stained cells were analyzed using FACSMelody cell sorter/analyzer (BD Biosciences), as described elsewhere ^21^. The FCS data files were analyzed using FlowJo data analysis software package.

### MTT-based cell proliferation assay

The PG cell suspension of passage 1 was prepared and dispensed at density of 2000 cells/well of a 96 well plate in 90 μl of media-1. The plates were incubated at 37°C, 5% CO_2_ for 24 hours. A serial dilution of H_2_O_2_ ranging from 0-1000μM in DMEM (Gibco) basal media is prepared in a separate 96 well plate. Subsequently, 10μl of each dilution is transferred respective to wells of plate with cells (0-100μM) and incubated for 48 hours. Subsequently, 10 μL MTT (5 mg/ml tetrazolium salt 3-[4,5-dimethylthiazol-2-yl]-2,5-diphenyltetrazolium bromide, Sigma-Aldrich) was added to each well and kept in a dark for 4 hours at 37°C. The MTT crystal were solubilized overnight in solubilization solution (10% SDS in 0.01N HCl) and absorbance was measured and at 570 nm and 660 nm was measured using Cytation 5 plate reader (BioTek) ^21^.

### *PSMA* gene promoter CpG methylation analysis

DNA samples 1μg obtained from SMG cells underwent bisulfite conversion using the Epitect Bisulfite Kit (Qiagen). Genomic DNA was denatured by sodium hydroxide treatment, followed by bisulfite conversion to selectively modify unmethylated cytosines to uracils while preserving methylated cytosines. The resulting bisulfite-converted DNA was purified using Epitect spin columns. Desulfonation was carried out to remove sulfonate groups, and after successive washes, the bisulfite-converted DNA was eluted with an appropriate buffer. Primers targeting the PSMA CpG islands from the bisulfite-converted DNA were designed using Meth Primer software ^69^. PCR amplification was performed using Q5 high-fidelity polymerase (NEB) to amplify the targeted regions. The resulting amplicons were subjected to Sanger sequencing. Sequencing data were analyzed using the web-based tool QUMA (QUantification tool for Methylation Analysis). QUMA facilitated bisulfite sequencing analysis of CpG methylation, providing essential data-processing functions and ensuring consistent quality control throughout the analysis process ^70^.

### Salivary single-cell RNA sequencing analysis

Adult C3H Mus musculus SMG single-cell RNA sequencing (scRNA-seq) data (GSM4546898) from Hauser et al. ^45^ was analyzed in R and R-Studio. Data analysis and visualization were performed using Seurat v5.0.1, tidyverse v2.0.0, and ggplot2 v3.4.4. Cells with fewer than 200 features, less than 1% of UMIs mapping to ribosomal genes, or greater than 10% UMIs mapping to mitochondrial genes were considered non-viable and excluded from analysis. Cells with greater than 2500 features were removed as an extra precaution against cell doublets. Additionally, cells with greater than 0.5% of UMIs mapping to hemoglobin genes were removed from analysis to exclude blood cells. Normalization and scaling were performed following Seurat’s default pipeline. 30 PCs from principal component analysis were computed to perform dimensionality reduction for clustering. Cell types were annotated based on protein and gene expression signatures cross-referenced through Human Protein Atlas immunostaining, Mouse Genome Informatics ^71^, Enrichr ^72^, Hauser et al. 2020 ^45^, and additional literature review. Murine analogs of human proteins were manually evaluated through cross-referencing Mouse Genome Informatics ^71^ and HUGO Gene Nomenclature Committee ^73^ databases.

### Sample preparation for salivary single-cell proteomics

Following FACS-based separation of viable progenitor-rich salivary single cells from SMG and PG, cells were sorted, and reactions were performed on a 384-well plate using the cellenONE system (Cellenion, France) ^42^. Lysis buffer was prepared with the concentration of 0.2% DDM (324355-1GM, Millipore, USA), 100 mM TEAB (T7408-500ML, Sigma-Aldrich, USA) and 2 ng/µl trypsin protease (90057, Thermo Scientific, USA). Prior to single-cell deposition, 1000 droplets (∼350 nl) of lysis buffer were dispensed into each well of 384-well plate. Single cells were then isolated and deposited into the wells containing lysis buffer. Plate was incubated on the heating deck inside the cellenONE at 37°C for 1 hour. Digestion was then quenched by adding 100 nl of 5% formic acid. Digested samples from each well were reconstituted in 4 µl of 0.1% formic acid containing 0.05x iRT peptides (Biognosys, Ki-3002-1).

### Liquid chromatography with tandem mass spectrometry (LC-MS/MS) data acquisition and analysis

Peptide samples were directly injected from the 384-well plate and separated on an analytical column (15 cm x 75 µm, C_18_ 1.7 µm, IonOpticks, AUR3-15075C18-CSI) using nanoElute 2 liquid chromatography system (Bruker Daltonics, Bremen, Germany). Solvent A (0.1% formic acid in water) and solvent B (0.1% formic acid in acetonitrile) were used to generate linear gradient over 38 min using a flow rate of 250 nl/min; 2-27% solvent B in 20 min, 27-40% solvent B in 5 min, 30-60% solvent B in 5 min, maintaining at 60-80% solvent B for 5 min, and 5% solvent B for 5 min. Separated peptides were analyzed using timsTOF SCP mass spectrometer (Bruker Daltonics, Bremen, Germany) in diaPASEF mode ^40^. Ions in the range of 400-1000 m/z were monitored with isolation window of 25 m/z and 8 PASEF scans per cycle along with 3 steps per PASEF scan.

The raw mass spectrometry data were analyzed using DIA-NN (version 1.8) using the following settings: protein inference=genes, quantification strategy=Robust LC (high accuracy) and cross-run normalization=global. UniProt human protein database (20,430 entries) were used for library-free analysis.

### Statistical analysis

All analyses were performed using GraphPad Prism 8.0. Differences were considered significant at p-value < 0.05. Data is displayed as mean±SEM unless otherwise stated.

## Author contributions

M.A., A.R.V., A.K., R.M. contributed to biobanking, data collection and analysis and preparation of the manuscript draft. D.G.M. and A.P. contributed to mass spectrometry data and analysis. R.C. and I.L. critically reviewed and assisted with manuscript preparation. J.J. and N.K. funded the study. N.K. conceptualized, designed and supervised the study and finalized the manuscript. All authors contributed to the drafting of the manuscript.

## Acknowledgements

Our heartfelt thanks go to all our donors for their invaluable contributions. We acknowledge the technical assistance provided by Geng Xian Shi, Wenmei Yang, Guruprasad Kalthur and Ishaq Viringipurampeer. Special appreciation to our team of Clinical Study Coordinators Stephanie Hafner, Matthew Dwarika, Miriam Anacker, Meaghan Rodgers, Elizabeth Starck and program coordinators Amanda Arnold and Betty Salerno.

## Supplementary Figure Legends

**Figure S1. The Impact of Donor Age and Duration from Collection-to-Processing on the Viability of Cells from Cryopreserved Tissue-Organoids. A)** Plot depicting viability percentages of bulk salivary cells post tissue-organoid dissociation across donor age (in years). **B)** Plot illustrating the percent viable salivary cells after tissue-organoid dissociation by duration of collection-to-processing of tissues (in hours).

**Figure S2. Analysis for Biomarker Expression in Male and Female Salivary Cells by FACS.** Plot showing percentage expression of various biomarkers (CD49f, PSMA, CD24, EpCAM, SSTR2, CD44, CD34) by flow cytometry in salivary cells of male and female donors (n=6).

**Figure S3. Comprehensive Immunophenotypic Analysis of Salivary Cells. A)** Salivary cells underwent immunophenotypic analysis using the antibody cocktail detailed in Figure 2A, with data analyzed using FlowJo. FACS data from six independent experiments were concatenated, normalized, and subjected to clustering for biomarker distribution analysis via t-SNE analysis. t-SNE plot showing expression of PSMA, EpCAM, CD24, and CD49f in salivary cells. Coloring correlates with expression intensity. Dark grey indicates high expression, while light grey signifies the absence of the cell marker expression. **B)** Barplot showing the expression of biomarkers by FACS across various passages. * indicate p-value < 0.05.

**Figure S4. Comparative Analysis of Biomarker Expression and Organoid Growth in Distinct Media Formulations. A)** Dynamic changes in biomarker expression profiles of unsorted (bulk) cells embedded in Matrigel and cultured in media-1 and media-2 across three sequential passages (P1-P3). **B)** Comparative analysis of biomarker expression profiles within the 3D OIC assay between media-1and media-2. **C)** Comparison between matched patients (n=5) based on organoids generated in media-1 and media-2, categorized by distinct phenotypes seeded. Statistical analyses are denoted within the graphs, where ‘ns’ signifies non-significant differences and * p-value <0.05 and **** p-value < 0.0001.

**Figure S5. Cell Output Analysis from Single Cells.** Plots showing cell output per starting single cells obtained from **A)** SMG **B-C)** PG cells cultured with or without collagen in media-1 or N2 media.

**Figure S6. Microscopic Analysis of 3D-Matrigel Organoids.** Brightfield images (4x and 10x) of 3D-matrigel organoids obtained from FACS-enriched (CD24+PSMA-, CD24+PSMA+, EpCAM+CD49f+, EpCAM+CD49f-, CD49f+, and CD49f-) of salivary cell subpopulations (Scale 1000µM and 200µM).

**Figure S7. Correlation of Biomarker Expression in Salivary Cells by FACS.** Plots showing the correlation between expression of CD49f, EpCAM and CD24 markers in **A)** PG and **B)** SMG cells.

**Figure S8. Biomarker Profiles in PG and SMG Cells across Passages.** Merged t-SNE clusters depicting the expression of CD49f, PSMA, CD24, EpCAM, SSTR2, CD34, CD44, and CD31/CD45 in **A)** PG and **B)** SMG cells across passages P0 and P1. Coloring corresponds to expression intensity, where red indicates high expression, and blue signifies the absence of the cell marker expression. **C)** Heatmap illustrating the expression of PSMA, CD49f, CD24, EpCAM and SSTR2 across passages P0 and P1.

**Figure S8. SGSPC-expressed Cytokeratins and their Pattern of Immunohistochemical Staining within Salivary Acinar Myoepithelial Cells (https://www.proteinatlas.org/).**

**Table S1. Single-cell Proteomics Data of Progenitor-Enriched Salivary Epithelial Cells.**

**Table S2. Post-translational Modifications of Proteins Identified by Single-Cell Proteomics Analysis of Progenitor-Enriched Salivary Epithelial Cells.**

**Table S3. Details of Antibodies Used for Immunohistochemistry.**

**Table S4. Details of Antibodies Used for FACS Analysis.**

