## Supplementary figures and images for "The Mayo Clinic Salivary Tissue-Organoid Biobanking: A Resource for Salivary Regeneration Research"

### Supplementary Figures

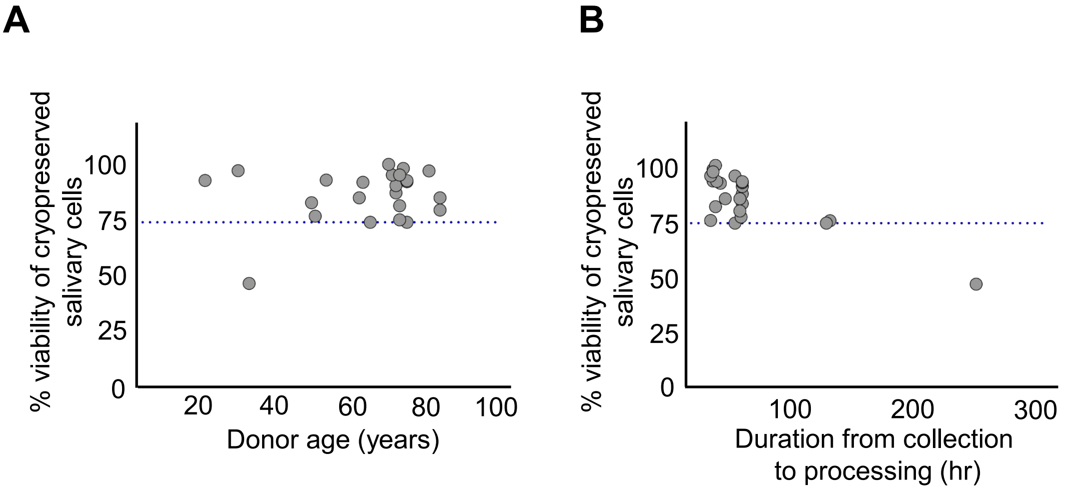

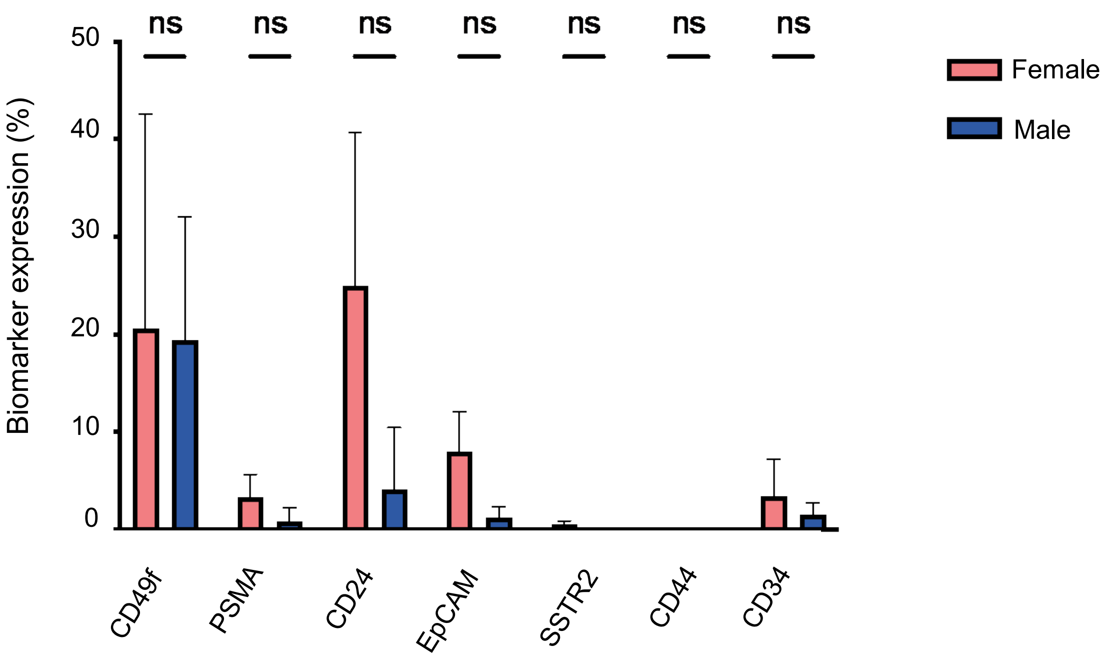

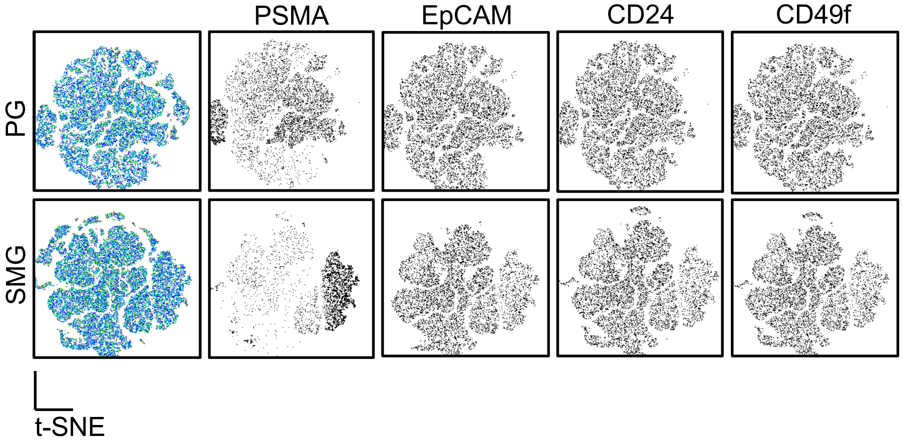

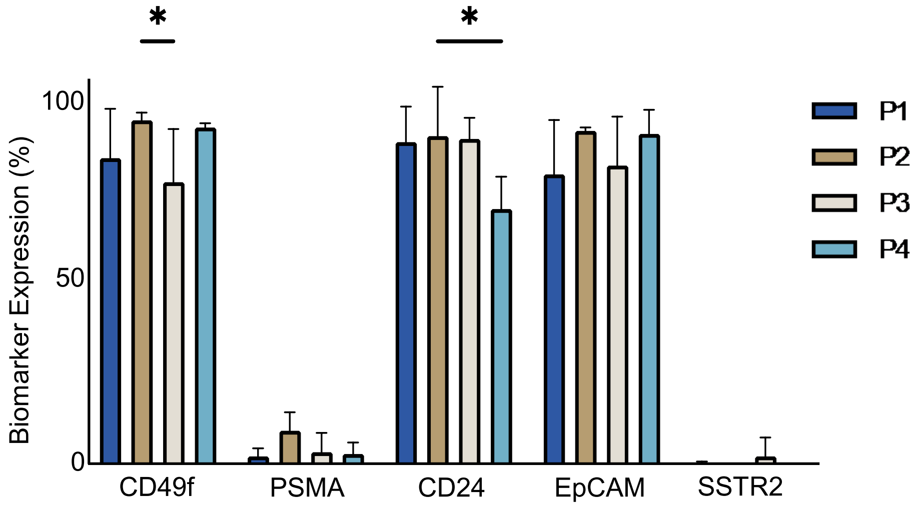

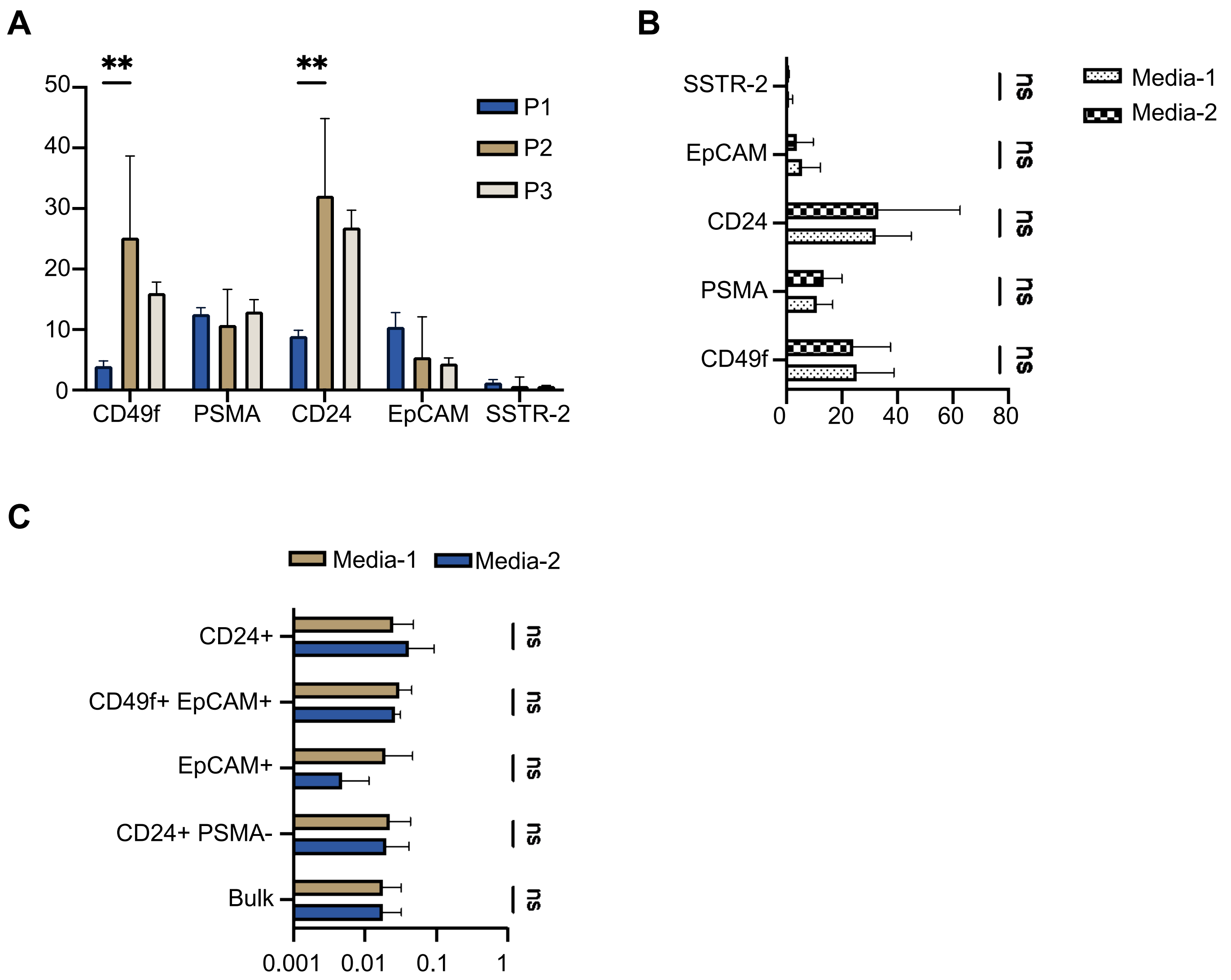

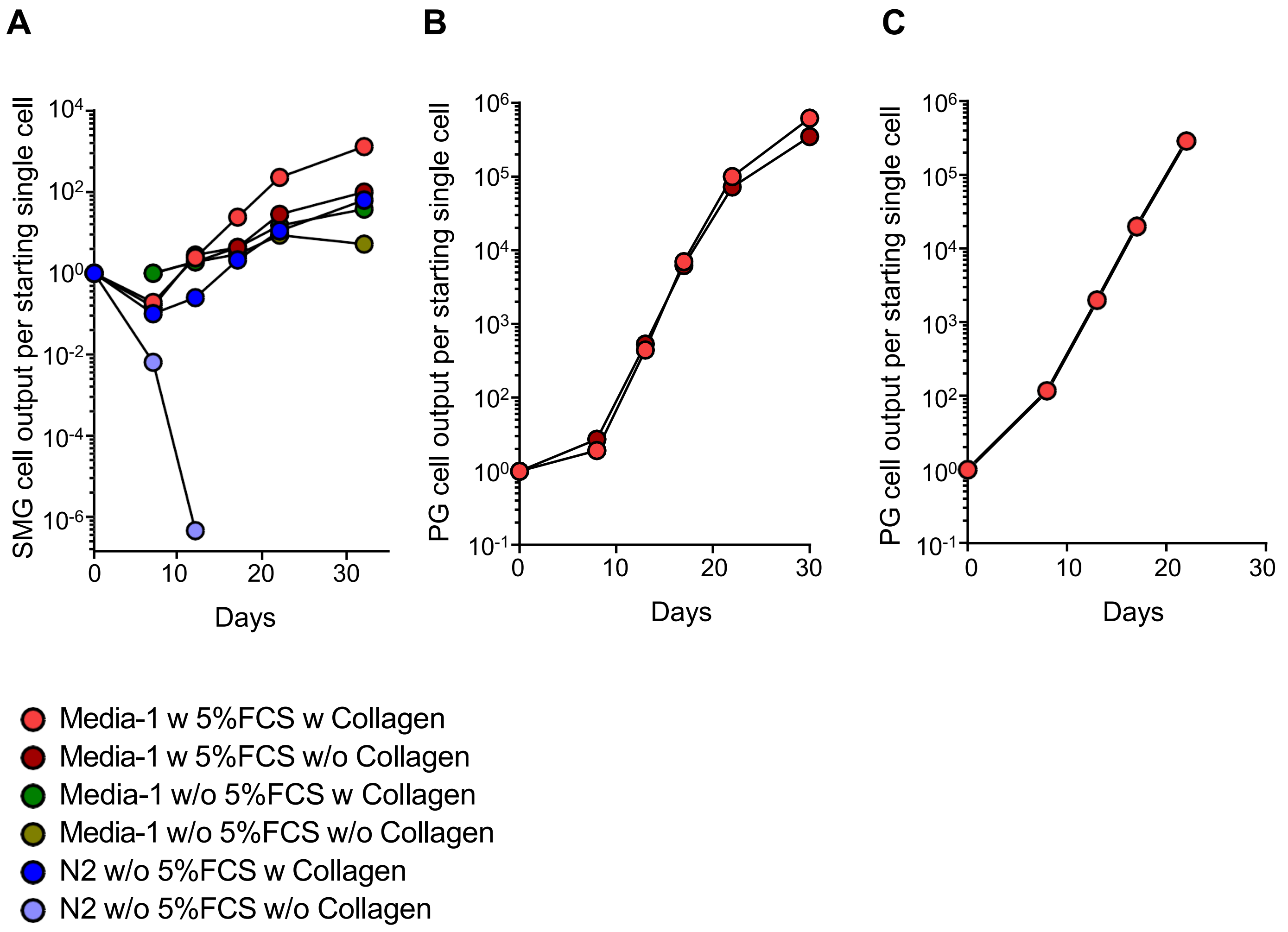

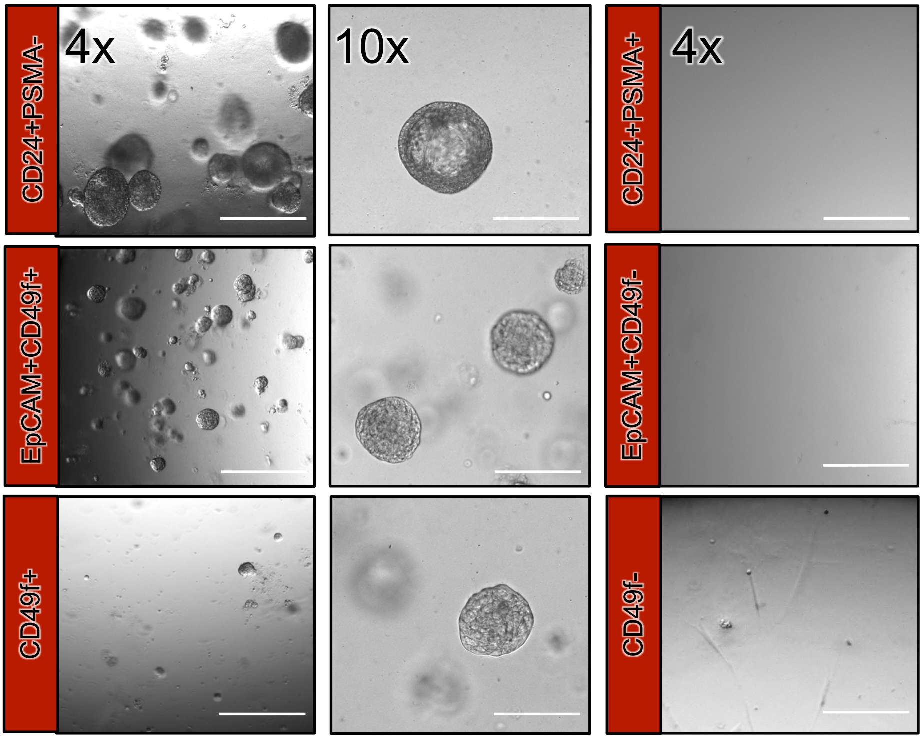

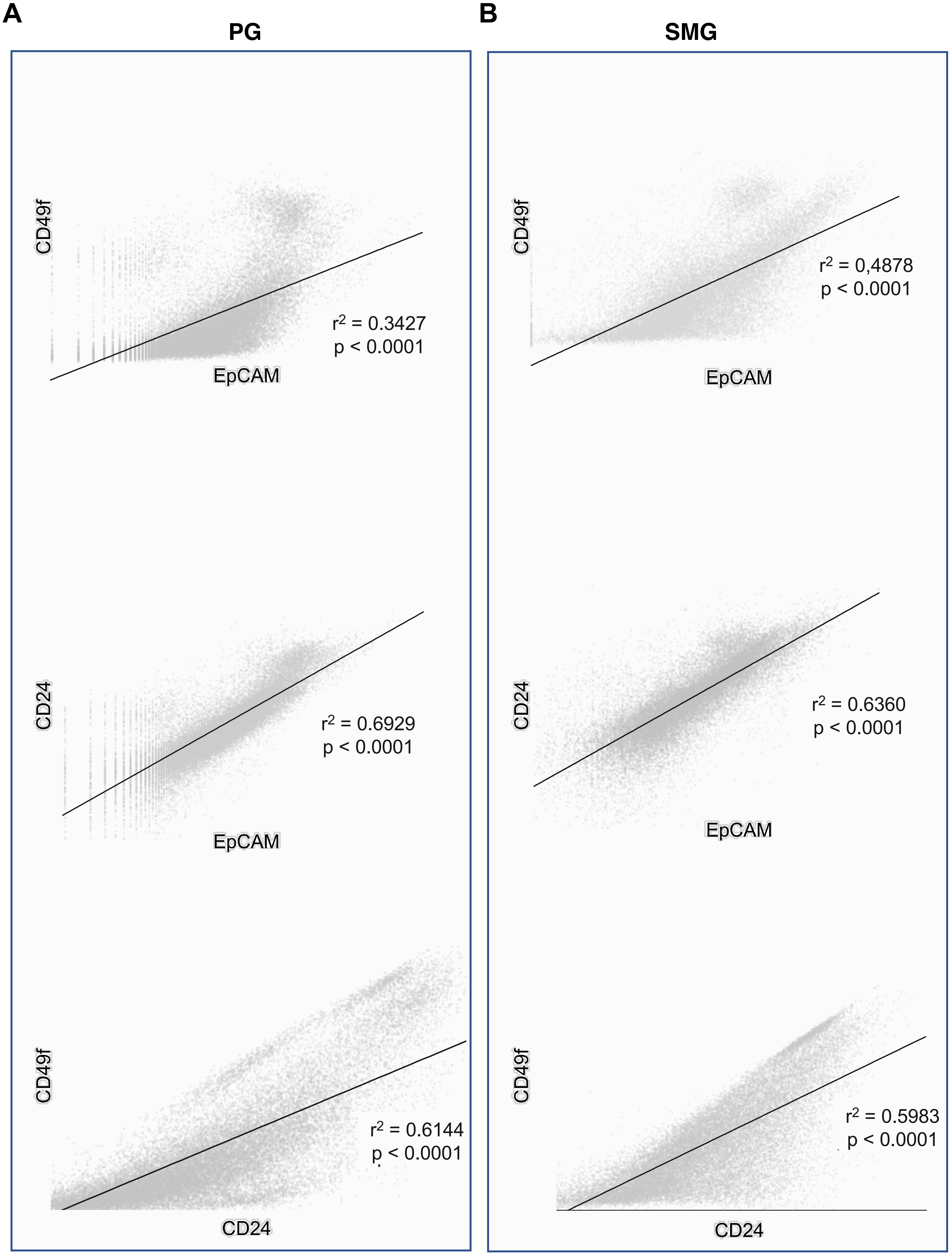

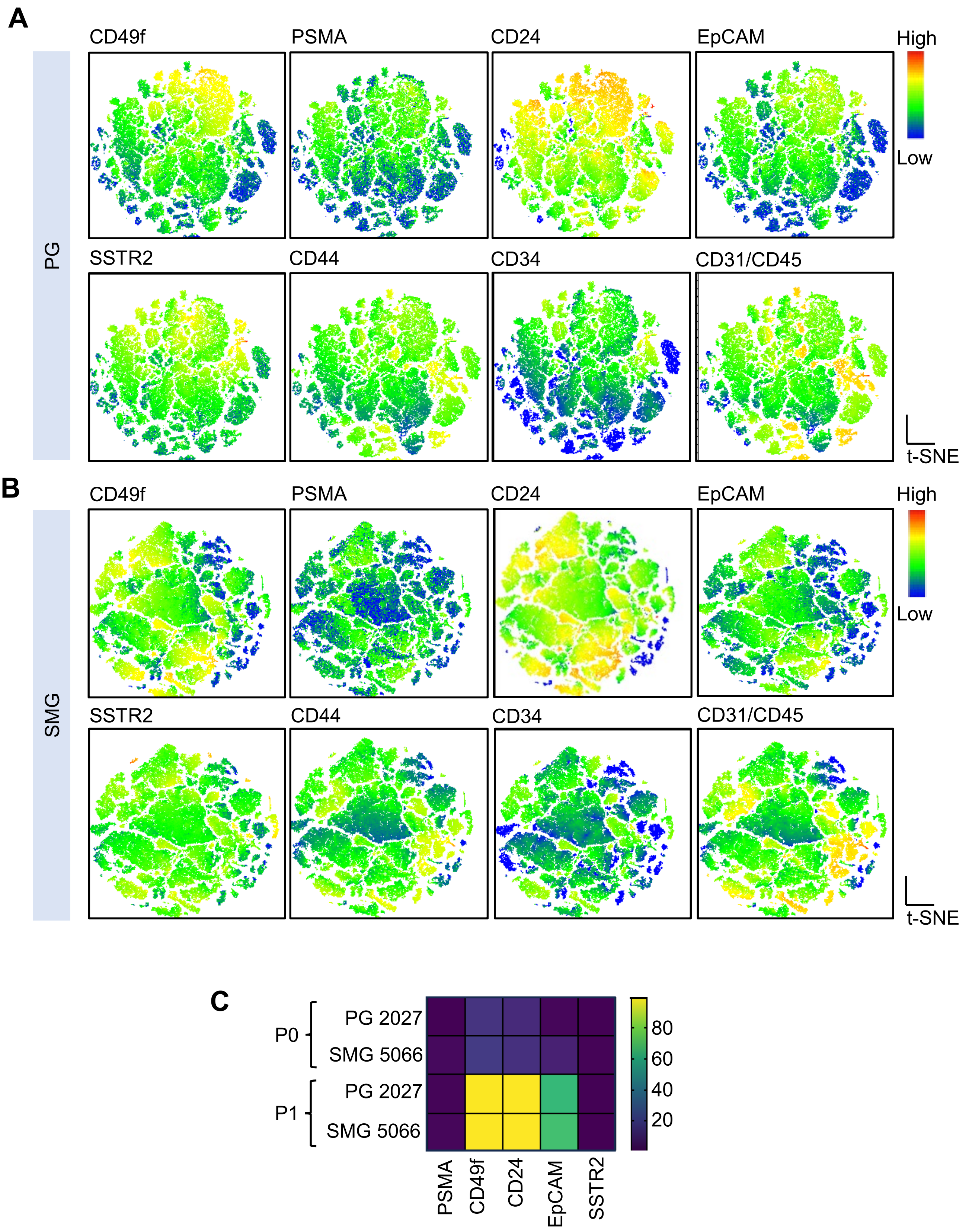

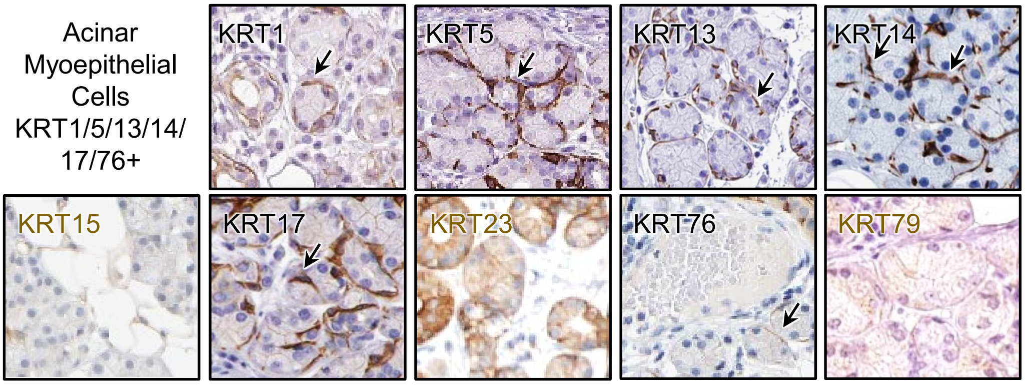
