## Supplementary material for "The Mayo Clinic Salivary Tissue-Organoid Biobanking: A Resource for Salivary Regeneration Research": Table S3

**Supplementary Table 1:** Details of antibodies used for immunohistochemistry

| Primary and secondary antibodies | Dilution | Clone ID | Catalog # | Vendor |
| --- | --- | --- | --- | --- |
| Anti-NKCC1 | 1:250 | NA | ab59791 | Abcam |
| Anti-Aquaporin-5 | 1:200 | NA | AB15858 | Millipore Sigma |
| Anti-Cytokeratin 5 | 1:500 | RCK105 | ab9021 | Abcam |
| Anti-Cytokeratin 7 | 1:250 | EP160Y | ab52635 | Abcam |
| Anti-Cytokeratin 8 | 1:250 | TROMA-1 | MABT329 | Millipore Sigma |
| Anti- Actin, Smooth Muscle | 1:250 | Ab-1(1A4) | MA1-06110 | Thermo Fisher Scientific |
| Alexa Fluor® 488 goat anti-rabbit IgG | 1:500 | NA | A-11008 | Thermo Fisher Scientific |
| Alexa Fluor® 594 goat anti-rabbit IgG | 1:500 | NA | A-11012 | Thermo Fisher Scientific |
