## Supplementary material for "The Mayo Clinic Salivary Tissue-Organoid Biobanking: A Resource for Salivary Regeneration Research": Table S4

**Supplementary table 2** – Details of the antibodies used for FACS analysis.

| Antibody | Antibody Clone | Antibody Source | Flourochrome |
| --- | --- | --- | --- |
| Anti-PSMA | LNI-I7 | BioLegend | PE-Cy7 |
| Anti-EpCAM | 9C4 | BioLegend | PerCP-Cy5.5 |
| Anti-SSTR2 | 402038 | R&D Systems | PE |
| Anti-CD34 | 561 | BioLegend | Alexa 488 |
| Anti-CD44 | 515 | BD Biosciences | BV510 |
| Anti-CD31 | WM59 | BioLegend | PacBlue |
| Anti-CD45 | HI30 | BioLegend | PacBlue |
| Anti-CD24 | ML5 | BioLegend | APC-Cy7 |
| Anti-CD49f | GoH3 | R&D Systems | APC |
